# *Klebsiella pneumoniae* disrupts vasodilation by targeting eNOS post translational modifications via the type VI secretion system and the capsule polysaccharide

**DOI:** 10.1101/2025.02.05.636584

**Authors:** Safi Rehman, Joana sa Pessoa, Charlotte Buckley, Rebecca Lancaster, Ciara Ross, Xun Zhang, John G. McCarron, Tim M. Curtis, Jose A. Bengoechea

## Abstract

Vasodilation is a crucial protective response to inflammation and infection. Endothelial cells control vasodilation through the bioavailability of eNOS-produced nitric oxide (NO), and the generation of endothelium-dependent hyperpolarization (EDH). Here, we demonstrate that *Klebsiella pneumoniae*, one of the most prevalent blood stream infection pathogens, inhibits agonist-induced vasodilation by blunting the NO-dependent pathway and attenuating the EDH pathway. The type VI secretion system (T6SS) effector VgrG4 licences the kinase PKCβ in an NLRX1-controlled mitochondria reactive oxygen species (mtROS)-dependent manner to phosphorylate the eNOS inhibitory site Thr^495^, effectively dampening eNOS activity. The capsule polysaccharide, on the other hand, limits the phosphorylation of the eNOS activation site Ser^1177^ by inducing the phosphatase PP2Ac upon activation of an EGF receptor-dependent pathway. VgrG4-induced mtROS attenuates the EDH pathway. Overall, this work reveals a new anti-host activity of the T6SS and illustrates how pathogens can control vascular biology by targeting eNOS post translational modifications.

## INTRODUCTION

Vascular endothelial cells are crucial to vascular biology, including the regulation of blood flow, maintenance of haemostatic balance, regulation of coagulation, control of vessel-wall permeability and the recruitment of leukocytes to sites of inflammation. Endothelium-governed vasodilation is a normal response to inflammation and infection to increase blood flow through affected areas and enabling increased delivery of immune cells for defence and repair. Not surprisingly, endothelial dysfunction, characterized by a shift towards reduced vasodilation and prothrombotic properties, is associated with the development of atherosclerosis, angiogenesis in cancer, vascular leakage, stroke, and infectious diseases^1^. Several viral infections induce endothelial dysfunction, being particularly dramatic in the case of haemorrhagic viruses such as Ebola, Lassa, and Marburg^2^. While few bacterial pathogens have been shown to infect endothelial cells^2^, there is a poor understanding of the effect of bacterial infections on vascular biology beyond the contribution of endothelial dysfunction to bacteria-induced sepsis^3^.

Vascular vasodilation is dependent on the bioavailability of endothelium-derived nitric oxide (NO)^4^, and, therefore, imbalances in the levels of NO cause endothelial dysfunction. Mechanistically, NO induces vasodilation upon diffusion into vascular smooth muscle cells where it activates soluble guanylyl cyclase (sGC) to form cyclic guanosine monophosphate (cGMP)^5^. NO is produced from L-arginine in a reaction catalysed by the constitutively expressed endothelial isoform of the enzyme nitric oxide synthase (eNOS)^5^. The activation of eNOS is strictly dependent on Ca^2+/^calmodulin^6,7^; and, therefore, an increase in the intracellular free Ca^2+^ concentration is necessary for endothelial NO production to trigger vasodilatation^7,8^. Apart from Ca^2+^, post translation modifications of eNOS, including phosphorylation, also regulate its activity^9,10^. Loss of eNOS function is associated with increased susceptibility to atherosclerosis, hypertension, thrombosis, sepsis, and stroke, reflecting the crucial role of NO and vasodilation in vascular biology. In addition to NO-dependent vasodilation, vascular endothelial cells of many vessels, such as small arteries, also regulate the vascular smooth muscle cell contractility through endothelium-dependent hyperpolarization (EDH)^11^. In this pathway, an increase in intracellular Ca^2+^ activates small (SKCa) and intermediate conductance (IKCa) Ca^2+-^ activated K^+^ channels, resulting in endothelial hyperpolarization^11^. The current produced subsequently spreads from the endothelium to the vascular smooth muscle through myoendothelial gap junctions located on endothelial projections to trigger vessel relaxation^11^.

Bloodstream infections (BSI) are one of the most lethal infections, with an estimated crude mortality rate of up to 30%. Furthermore, BSI caused by antibiotic resistant bacteria are associated with increased mortality, and ICU admission compared to BSI caused by susceptible bacteria^12–16^. Of particular concern are the infections caused by *Klebsiella pneumoniae,* the second most prevalent Gram-negative pathogen in BSI, with a mortality rate of up to 79%^12,16,17^, owing to the increasing isolation of multidrug resistant strains globally. Despite the clinical relevance of *Klebsiella*-induced BSI, there is a gap in our knowledge of the interface between *K. pneumoniae* and the vasculature. Intriguingly, *K. pneumoniae*-derived pneumosepsis shares several hallmarks with acute respiratory distress syndrome (ARDS), characterized by damage to the capillary endothelium and alveolar epithelium, as well as fluid accumulation in the alveolar space, leading to alveolar oedema. Notably, clinical studies have found an association between hypertension and *K. pneumoniae* infection^18^, and in vivo work showed limited vasodilation in arteries of mice infected with *K. pneumoniae*^18^. Altogether, this evidence suggests a crucial role of altered endothelial function in *K. pneumoniae* pathophysiology.

Here, we provide mechanistic insights into the interface between *K. pneumoniae* and blood vessels by leveraging a research platform that includes an ex vivo blood vessel model and human primary endothelial cells. This platform has allowed us to examine and quantify functional responses upon infection and identify signalling pathways targeted by *K. pneumoniae* that affect vascular physiology. We uncover that *K. pneumoniae* exerts a profound effect on the vasculature, resulting in the inhibition of agonist-induced vasodilation. We demonstrate that *K. pneumoniae*-induced reduction of NO bioavailability is mediated by the trans-kingdom type VI secretion (T6SS) effector VgrG4 and the capsule polysaccharide. VgrG4 triggers the phosphorylation of the eNOS inhibitory site Thr^495^ via activation of the kinase PKCβ in a mitochondria reactive oxygen species (mtROS)-dependent manner which occurs after the activation of the mitochondria located innate receptor NLRX1. The capsule polysaccharide, on the other hand, limits the phosphorylation of the eNOS Ser^1177^ governing eNOS activity via the phosphatase PP2Ac upon activation of an EGF receptor (EGFR)-dependent pathway. Collectively, these findings demonstrate that *K. pneumoniae* causes vascular endothelial dysfunction by targeting eNOS, unveiling previously unknow signalling cascades exploited by a pathogen to control vascular biology.

## RESULTS

### *K. pneumoniae* inhibits agonist-induced vasodilation

To investigate the effect of infections on the vasculature, we developed an ex vivo blood vessel model in which rat mesenteric arteries were mounted in a pressure myograph system. This set-up enabled us to quantify changes in vasoconstriction and vasodilation in real-time in response to various agonists and infections, with the outer diameter of the vessel serving as the primary measurement (see material and methods and Fig 1A). In control vessels, phenylephrine (PE) induced constriction, while the subsequent addition of acetylcholine (ACh), which activates endothelial M3-muscarinic receptors^19–21^, resulted in vasodilation (Fig 1B and movie S1). 100 nM ACh induced transient vasodilation (Fig 1B and movie S1), while at 1 μM, it restored the basal diameter of the vessel (Fig 1B and movie S2). As reported previously for these vessels^11,22^, inhibition of NO synthesis with L-NAME resulted in transient ACh-induced vasodilation, reaching the vessel’s basal diameter (Fig S1A). In turn, inhibition of the EDH pathway with the IKCa channel inhibitor TRAM-34 and the SKCa channel inhibitor apamin resulted in small, sustained increases in vasodilation with increasing concentrations of ACh. However, these increases never reached the vessel’s basal diameter (Fig S1B). The combination of the three inhibitors resulted in no ACh-induced vasodilation (Fig S1C), demonstrating that ACh-induced vasodilation in these vessels relies entirely on the NO and EDH signalling pathways.

**Figure 1.**
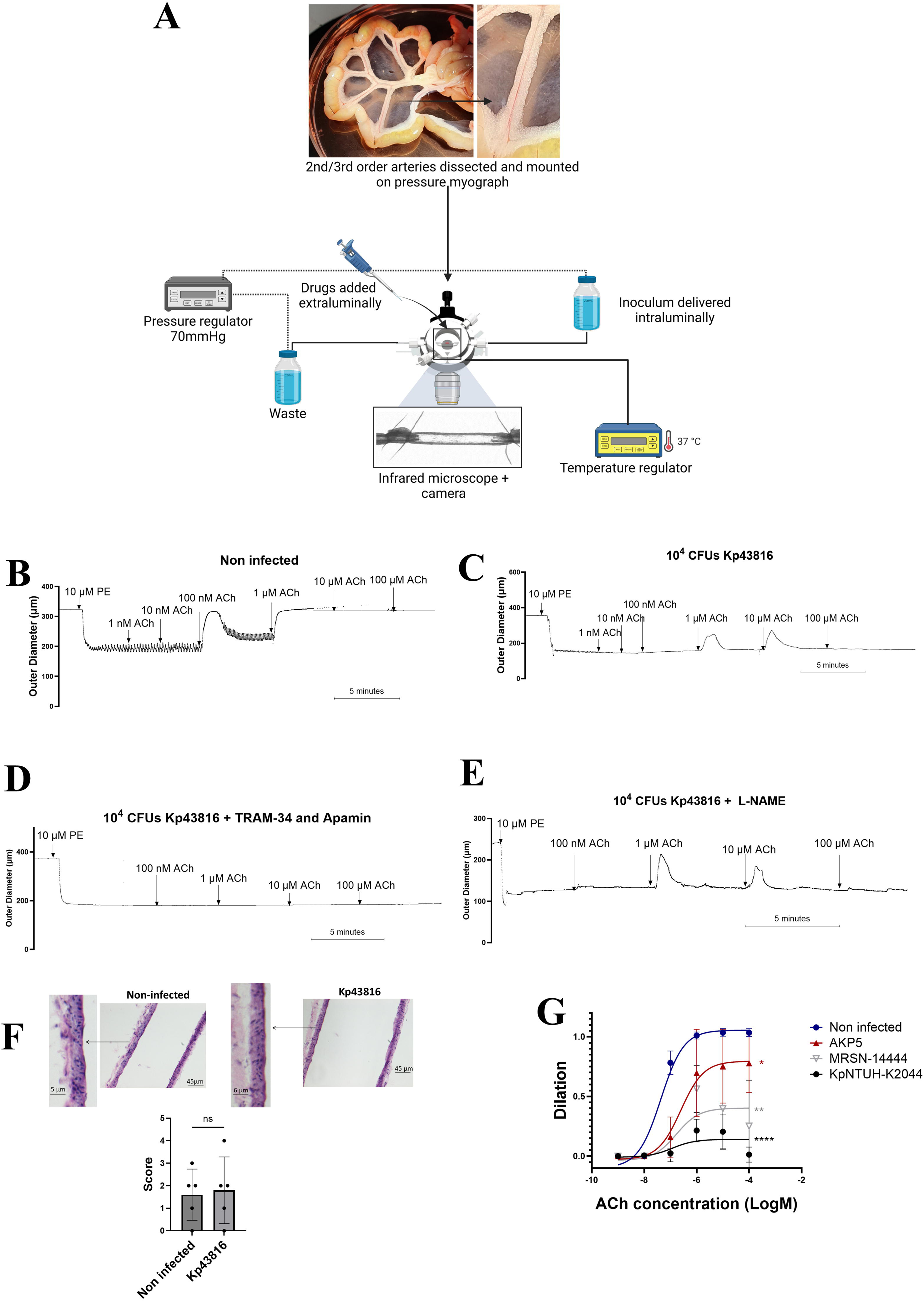
*K. pneumoniae* inhibits ACh-induced vasodilation. (A). Schematic model of the ex vivo blood vessel model. (B). Representative trace of ACh-induced vasodilation in PE-constricted vessels. (C). Representative trace of Kp43816-triggered inhibition of ACh-induced vasodilation in PE-constricted vessels. (D). Representative trace of the effect of EDH pathway inhibitors TRAM-34 and apamin on Kp43816-triggered inhibition of ACh-induced vasodilation in PE-constricted vessels. (E). Representative trace of the lack of effect of the NO pathway inhibitor L-NAME on Kp43816-triggered inhibition of ACh-induced vasodilation in PE-constricted vessels. (F). Representative images of hematoxilin-eosine stained non-infected and infected vessels and scoring of blood vessel pathology. (G). Dose response of ACh-induced peak dilation in non-infected and infected vessels with strains AKP5, MRSN-14444, and NTUH-K2044. In B, C, D and E the traces are representative of six vessels from three rats. In F, images are representative of five vessels from five different rats. Histopathology quantification is shown as mean ± SD and analysed with Mann-Whitney t test. ns, p > 0.05. In G, peak dilation is shown as mean ± SD from six vessels from three rats per group and analysed with two-way ANOVA and Dunnet’s multiple group comparison correction. The asterisks indicate p < 0.05 (*), < 0.01 (**), or <0.0001 (****).

To ascertain whether *Klebsiella* infection affects vascular physiology, mesenteric arteries were infected with *K. pneumoniae* strain ATCC43816 (hereafter Kp43816). This strain encodes all the loci found in those strains associated with invasive community-acquired infections^23^. Intraluminal infection with 10^4^ CFUs Kp43816 neither affected the outer diameter of the vessel (Fig S1D) nor PE-induced vasoconstriction (Fig S1E). In contrast, Kp43816 inhibited ACh-induced vasodilation (Fig 1C, Fig S1F, movie S3 and movie S4. 1-log less bacteria also inhibited ACh-triggered vasodilation (Fig S1F and Fig S1G). TRAM-34 and apamin (Fig 1D) but not L-NAME (Fig 1E) abrogated the residual ACh-induced vasodilation in Kp43816-infected vessels suggesting that *K. pneumoniae* abolishes NO-dependent vasodilation. In addition, EDH-mediated vasodilation was significantly diminished when compared with control vessels. Chemical complementation using the NO donor sodium nitroprusside (SNP) restored vasodilation in infected vessels (Fig S1H). The fact that SNP-induced vasodilation was similar in control and infected vessels illustrates that the function of the vascular smooth muscle in response to NO was unaffected by intraluminal *Klebsiella*. Further demonstrating that infection did not result in overt vascular damage, histopathology analysis showed that infection did not affect the endothelium or smooth muscle layers (Fig 1F). 10^4^ CFUs UV-killed bacteria or 10^4^ latex beads similar in size to *Klebsiella* did not impact ACh-dependent vasodilation (Fig S1I), thus ruling out the possibility that the mere presence of bacteria has a detrimental effect on ACh-induced vasodilation. Strain NTUH-K2044^24^, a clinical isolate used to probe *K. pneumoniae* metastatic strains found in Asia, and the clinical isolates AKP5 and MRSN-14444 also inhibited ACh-induced vasodilation (Fig 1G). This reveals that *Klebsiella* inhibition of ACh-mediated vasodilation is not strain dependent.

Combined these data demonstrate that live *K. pneumoniae* abrogates ACh-induced vasodilation in mesenteric arteries without damaging the endothelial cell layer and perturbing the function of the vascular smooth muscle. Further experiments using inhibitors targeting the specific pathways governing vasodilation in these vessels uncovered that *K. pneumoniae* blunts the NO pathway and attenuates the EDH pathway to inhibit vasodilation.

### The trans-kingdom T6SS effector VgrG4 mediates *K. pneumoniae* inhibition of vasodilation

We next sought to identify the *Klebsiella* factor(s) responsible for the inhibition of vasodilation. Considering the crucial role of the CPS on *Klebsiella*-host interactions^25^, we investigated a *cps* mutant to assess whether the CPS is involved in the inhibition of ACh-induced vasodilation. However, the *cps* mutant still abrogated ACh-induced vasodilation (Fig 2A and Fig S2A). Recently, we have uncovered the role of the T6SS as an immune evasin^26,27^. Consequently, we asked whether the T6SS was responsible for the *Klebsiella*-induced inhibition of vasodilation. In contrast to Kp43816-infected vessels, ACh-induced relaxation was not abrogated in *tssB* mutant infected vessels (Fig 2A and Fig S2B). TssB is a core component of the sheath section of the T6SS apparatus, and the T6SS is not assembled in a *tssB* mutant^28–31^. The T6SS also mediates NTUH-K2044-induced inhibition of ACh-dependent vasodilation because a *clpV* mutant did not abrogate ACh-triggered vasodilation (Fig S2C). A *clpV* mutant cannot assemble a functional T6SS^28–31^.

**Figure 2.**
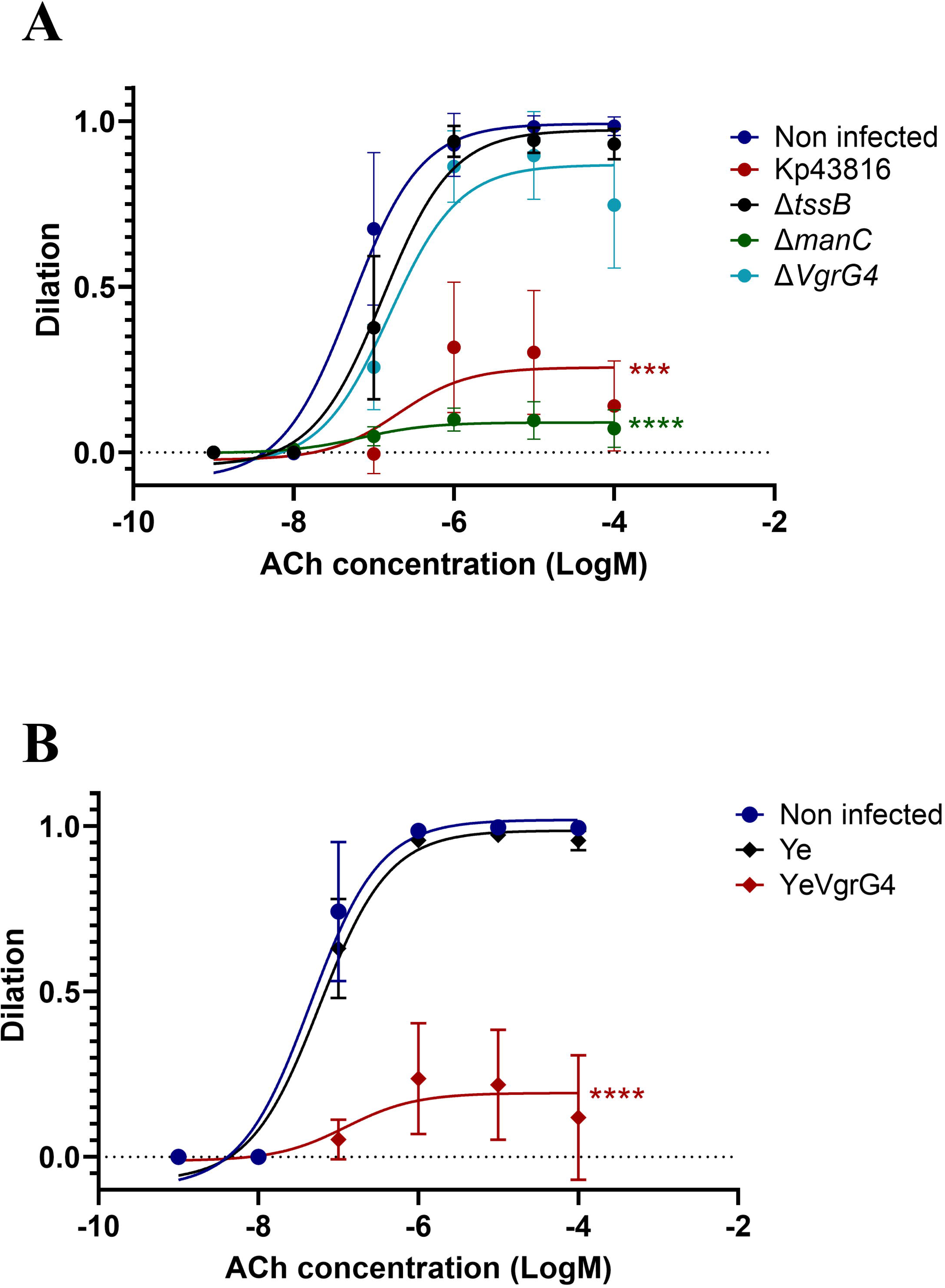
T6SS effector VgrG4 inhibits ACh-induced vasodilation. (A). Concentration response of ACh-induced peak dilation in non-infected and infected vessels with strains Kp43186, *cps* mutant (Δ*manC*; 43816-Δ*manC*) and the T6SS mutants tssB (Δ*tssB*; 43816-Δ*tssB* and *vgrG4* (Δ*vgrG4*; 43816-Δ*vgrG4*). (B). Concentration response of ACh-induced peak dilation in non-infected and infected vessels with strains Ye and YeVgrG4. In A and B, peak dilation is shown as mean ± SD from six vessels from three rats per group and analysed with two-way ANOVA and Dunnet’s multiple group comparison correction. The asterisks indicate p < 0.001 (***), or <0.0001 (****).

Next, we aimed to identify the T6SS effector mediating the inhibition of ACh-triggered vasodilation. We initially focused on the trans-kingdom effector VgrG4^26,32^ because we have recently demonstrated that *Klebsiella* injects this effector into mammalian cells in a T6SS-dependent manner^26^. The *vgrG4* mutant did not impair ACh-triggered vasodilation (Fig 2A and Fig S2D). Because VgrG proteins are also known to act as cargo for other T6SS effectors^33^, the possibility exists that the inhibition of vasodilation could be mediated by other T6SS effector(s) delivered by VgrG4. To exclude this possibility, we took advantage of a *Yersinia* toolbox that enables the study of the cellular effects of single bacterial effectors through *Yersinia* type 3 secretion system (Ysc)-T3SS-mediated injection into eukaryotic cells^34^. We previously confirmed the secretion of VgrG4 in conditions in which the T3SS is active^26^. Intraluminal infection of vessels with the *Y. enterocolitica* control strain (hereafter Ye) did not affect ACh-induced vasodilation (Fig 2B and Fig S2E). In contrast, *Y. enterocolitica* encoding *vgrG4* (hereafter YeVgrG4) abrogated ACh-induced vasodilation (Fig 2B and Fig S2F), demonstrating that *K. pneumoniae* T6SS effector VgrG4 is necessary and sufficient to inhibit ACh-triggered vasodilation.

### *K. pneumoniae* does not impair endothelial Ca^2+^ signalling

We next sought to provide mechanistic insights into how *Klebsiella* inhibits vasodilation in a T6SS VgrG4-dependent manner. The activation of the NO and EDH pathways in the vascular endothelium is strictly dependent on Ca^2+-^signalling^6,7,11^. In fact, evidence establishes a linear relationship between population-level endothelial Ca^2+^ responses and vasodilation^35,36^. Therefore, *Klebsiella*-mediated inhibition of both pathways could be explained by impaired Ca^2+^ responses in endothelial cells upon infection. To start addressing this question, we infected immortalized human lung microvascular endothelial cells (HULEC-5a, hereafter HULEC) and assessed Ca^2+^ responses using Fura-2 dye. We used histamine instead of ACh for these experiments because HULEC cells did not respond to ACh (Fig S3A) due to the lack of expression of the M3 muscarinic receptor^37^. Kp43816 did not inhibit histamine induced Ca^2+^ responses (Fig 3A). In fact, at low concentrations of histamine, infection increased the Ca^2+^ responses. (Fig 3A). These observations suggest that *Klebsiella* may not affect endothelial Ca^2+^ responses.

**Figure 3.**
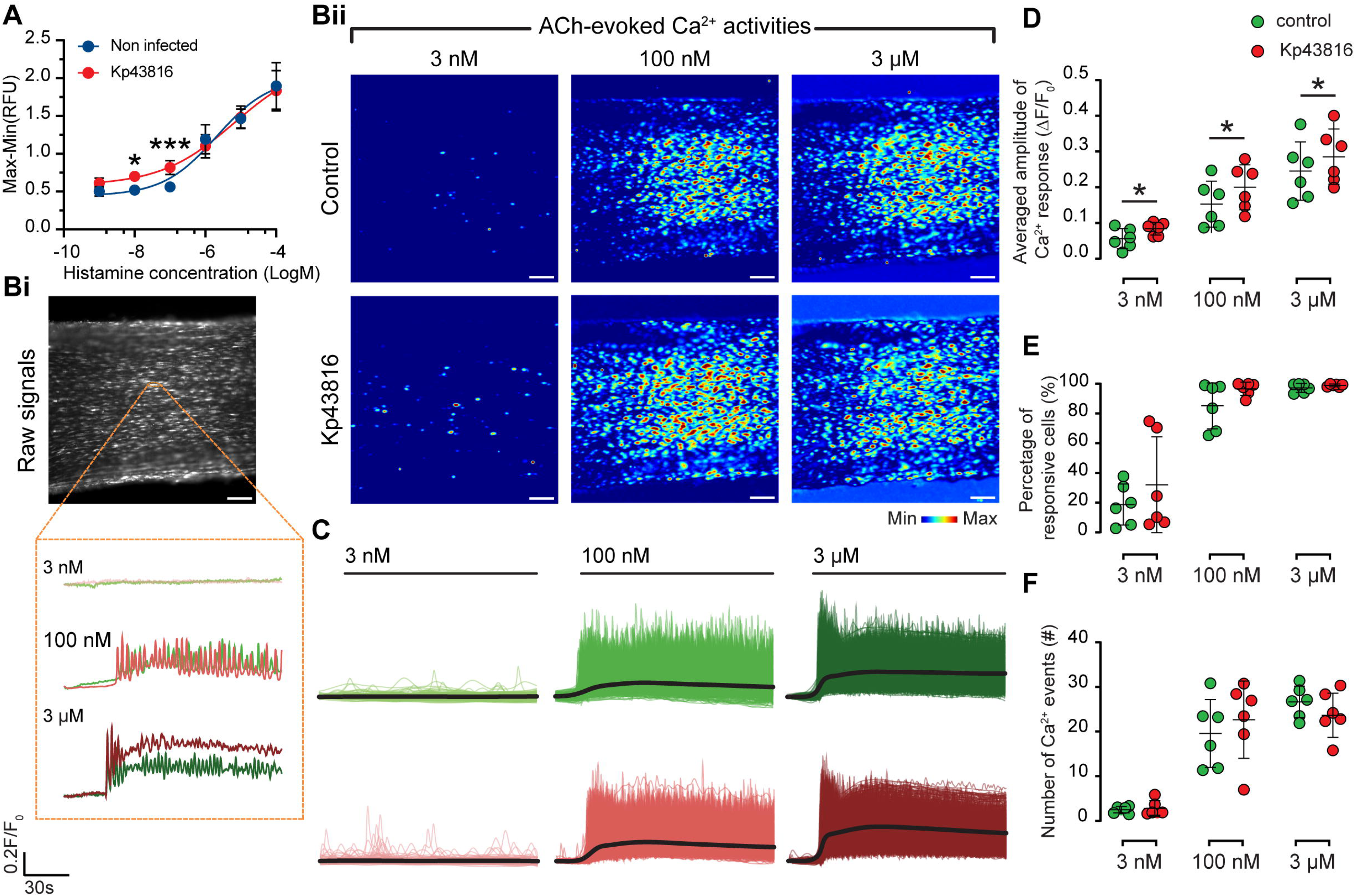
*K. pneumoniae* does not impair endothelial Ca^2+^ signalling. (A). Concentration response of histamine-induced intracellular Ca^2+^ in non-infected and Kp43816-infected HULEC cells. (B). Averaged Ca^2+^ signal in response to ACh in non-infected and kp43816-infected vessels. (C). Mean amplitude of Ca^2+^ response of a single cell to ACh in non-infected and kp43816-infected vessels. (D). Frequency of Ca^2+^ oscillation evoked by ACh in non-infected and kp43816-infected vessels. (E). Mean percentage of active cells evoked by ACh in non-infected and kp43816-infected vessels. In A, data is shown as mean ± SD from three independent experiments in sextuplicate. In C, D and E, each dot represents the data from one vessel before and after infection, and the mean is indicated. Data were analysed with two-way ANOVA and Dunnet’s multiple group comparison correction. The asterisks indicate p < 0.05 (*). <0.001 (***).

To determine whether *Klebsiella* impairs Ca^2+^ signalling in our experimental model, we examined endothelial Ca^2+^ activity in intact blood vessels using Cal-520 coupled with high spatiotemporal resolution, wide-field single photon imaging in fields of ∼1,000 cells (Fig 3). In these experiments, we quantified the responses of individual endothelial cells across the entire population (Fig 3B) to three concentrations of ACh (3, nM, 100 nM and 3 µM) over a 200 s recording period. The averaged response amplitude for individual cells (Fig 3C and Fig 3D), the percentage of cells responding (Fig 3E), and the frequency (number) of single cell Ca^2+^ events (Fig 3F) for each vessel was measured. Infection did not evoke any Ca^2+^ response (movie S5). In control vessels, ACh produced a concentration-dependent, distributed and heterogeneous response across the field of endothelial cells (Fig 3B and movie S4), as previously described^35^. Infection of the endothelium did not itself evoke a Ca^2+^ response in endothelial cells (movie S4). Contrary to expectations, infection did not diminish the Ca^2+^ response to ACh. In fact, the amplitude of the ACh-induced Ca^2+^ signal was greater in both individual cells and the overall population in infected vessels (Fig 3C and Fig 3D; movie S5). Furthermore, quantification of the number of cells responding to ACh (Fig 3E) and the number of Ca^2+^ events in each cell (Fig 3F) illustrated that Kp43816 infection did not reduce the ACh-induced Ca^2+^ response. Control experiments confirmed that the response to ACh in non-infected and infected vessels was mediated by muscarinic M3 receptors (inhibited by 4-DAMP; 1 μM) (Fig S3B). These findings are consistent with previous studies demonstrating the role of these receptors in endothelial cells responses to ACh^19–21^.

Altogether, this evidence shows that ACh-evoked Ca^2+^ responses are increased in *K. pneumoniae*-infected vessels, suggesting that *K. pneumoniae*-induced blockade of vasodilation cannot be attributed to impaired endothelial Ca^2+^ responses.

### *K. pneumoniae* controls post translational modifications of eNOS to blunt its function

Evidence so far demonstrates that *Klebsiella* blunts the NO pathway, with vasodilation restored by employing an NO donor. The fact that *Klebsiella* infection does not impair Ca^2+^ responses, an upstream event required for eNOS activation^6,7^, prompted us to test whether *Klebsiella* may affect the protein expression of eNOS in an VgrG4-dependent manner to abrogate the NO pathway. We infected HULEC cells, and eNOS levels were assessed by immunoblotting. Infection did not affect the levels of eNOS (Fig 4A). Similar results were obtained when we infected primary human pulmonary microvascular endothelial cells (HPMEC) (Fig S4A). We next considered whether VgrG4 may target eNOS post translational modifications that regulate its activity^9,10^. The phosphorylation of Ser^1177^ results in activation of the enzyme^38,39^ whereas the phosphorylation of Thr^495^, located in the eNOS calmodulin binding site, inhibits eNOS activity^38,39^. eNOS agonists induce the phosphorylation of Ser^1177^ whereas drugs that inhibit eNOS activity induce the phosphorylation of Thr^495^ ^38,39^. Moreover, the phosphorylation of Thr^495^ is sufficient to inhibit eNOS activity independently of the phosphorylation status of Ser^1177^ ^39^. We initially investigated whether *Klebsiella* targets the phosphorylation of Ser^1177^. Immunoblot analyses showed that Kp43816 did not induce the phosphorylation of Ser^1177^ unlike bradykinin, histamine and ionomycin, three agonists known to activate eNOS (Fig 4B). Notably, Kp43816 inhibited the agonist-induced phosphorylation of Ser^1177^ (Fig 4B), suggesting that *Klebsiella* limits the phosphorylation of the activation site of eNOS. Control experiments confirmed that pre-incubation of the agonists with *Klebsiella* did not affect their capacity to induce the phosphorylation of Ser^1177^ (Fig S4B). We then assessed whether the T6SS effector VgrG4 mediates *Klebsiella*-controlled inhibition of Ser^1177^ phosphorylation. However, in cells infected with either the *tssB* mutant (Fig 4C) or YeVgrG4 (Fig 4D) we did not observe any decrease in the agonist-induced Ser^1177^ phosphorylation, indicating that the T6SS does not mediate the inhibition of Ser^1177^ phosphorylation. In contrast, agonist-induced Ser^1177^ phosphorylation was not inhibited in cells infected with the *cps* mutant (Fig 4E), revealing that *Klebsiella*-mediated inhibition of Ser^1177^ phosphorylation is CPS dependent.

**Figure 4.**
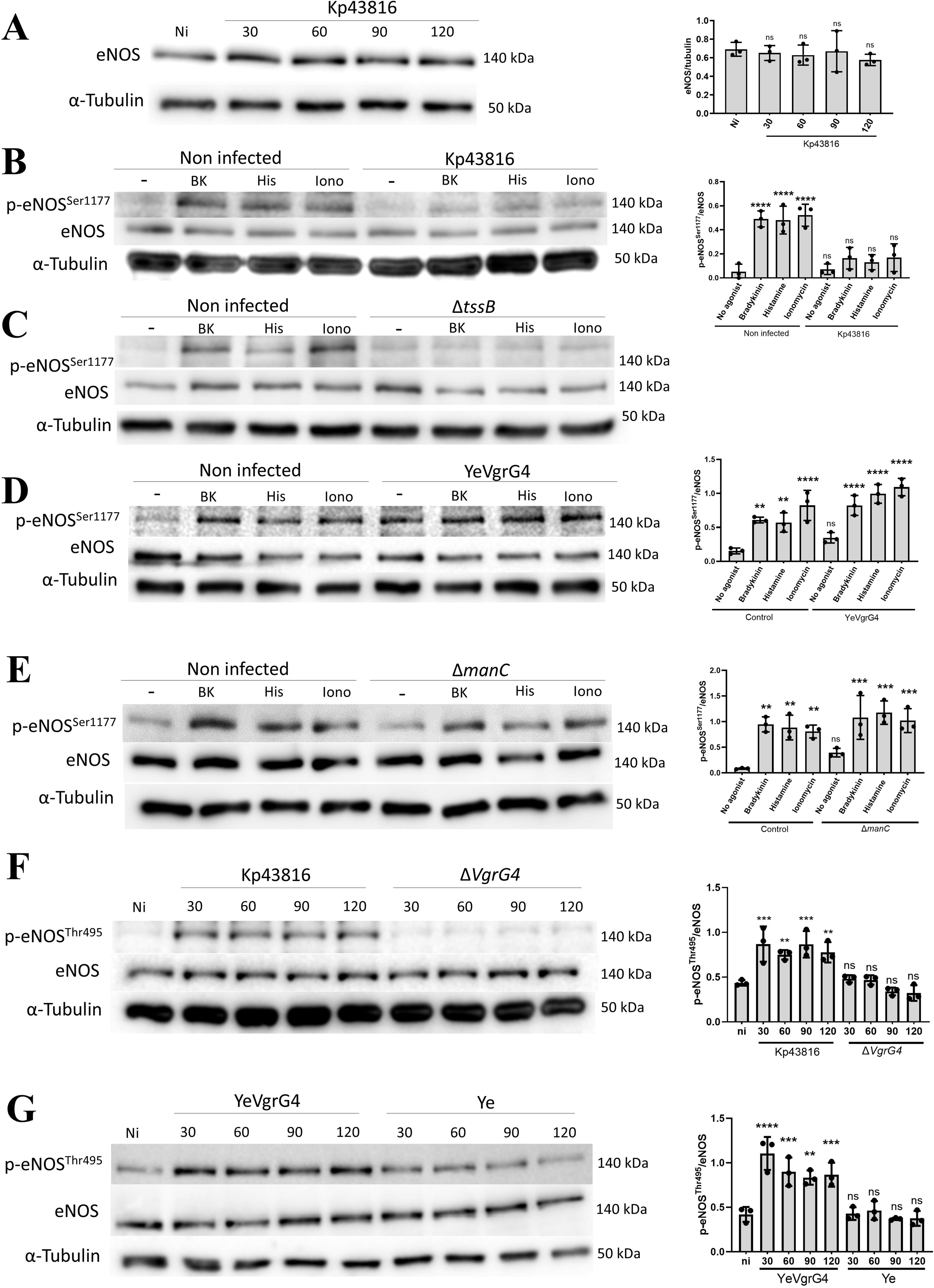
*K. pneumoniae* targets eNOS post translational modifications in a T6SS effector VgrG4 and capsule-dependent manner. (A). Immunoblot analysis of eNOS and tubulin levels in infected HULEC cells for the indicated time points. Densitometry analysis of blots (n = 3) representing the ratio of eNOS versus tubulin. (B). Immunoblot analysis of phosphorylated eNOS Ser^1177^, eNOS and tubulin levels in HULEC cells treated for 5 min with 1 μM bradykinin (Bk), histamine (His), and ionomycin (Iono). When indicated, cells were infected with Kp431816 for 120 min before adding the agonists. Densitometry analysis of blots (n = 3) representing the ratio of phosphorylated eNOS Ser^1177^ versus eNOS. (C). Immunoblot analysis of phosphorylated eNOS Ser^1177^, eNOS and tubulin levels in HULEC cells treated for 2 min with 1 μM bradykinin (Bk), histamine (His), and ionomycin (Iono). When indicated, cells were infected with the T6SS mutant *tssB* (Δ*tssB*; 43816-Δ*tssB*) for 120 min before adding the agonists. Densitometry analysis of blots (n = 3) representing the ratio of phosphorylated eNOS Ser^1177^ versus eNOS. (D). Immunoblot analysis of phosphorylated eNOS Ser^1177^, eNOS and tubulin levels in HULEC cells treated for 2 min with 1 μM bradykinin (Bk), histamine (His), and ionomycin (Iono). When indicated, cells were infected with YeVgrG4 for 120 min before adding the agonists. Densitometry analysis of blots (n = 3) representing the ratio of phosphorylated eNOS Ser^1177^ versus eNOS. (E). Immunoblot analysis of phosphorylated eNOS Ser^1177^, eNOS and tubulin levels in HULEC cells treated for 2 min with 1 μM bradykinin (Bk), histamine (His), and ionomycin (Iono). When indicated, cells were infected with the *cps* mutant (Δ*manC*; 43816-Δ*manC*) for 120 min before adding the agonists. Densitometry analysis of blots (n = 3) representing the ratio of phosphorylated eNOS Ser^1177^ versus eNOS. (F). Immunoblot analysis of phosphorylated eNOS Thr^495^, eNOS and tubulin levels in HULEC cells infected with Kp43816 and *vgrG4* mutant (Δ*vgrG4*; Kp43816-Δ*vgrG4*) for the indicated time points. Densitometry analysis of blots (n = 3) representing the ratio of phosphorylated eNOS Thr^495^ versus eNOS. (G). Immunoblot analysis of phosphorylated eNOS Thr^495^, eNOS and tubulin levels in HULEC cells infected with Ye and YeVgrG4 for the indicated time points. Densitometry analysis of blots (n = 3) representing the ratio of phosphorylated eNOS Thr^495^ versus eNOS. Blots are representative of three independent experiments. In all panels, data were compared against the non-infected control using one-way ANOVA and Tukey’s multiple group comparison correction. The asterisks indicate p <0.01 (**). <0.001 (***), or <0.0001 (****). ns, not significant.

We next questioned whether *Klebsiella* targets the phosphorylation of inhibitory site Thr^495^. Kp43816 induced the phosphorylation of Thr^495^ in both HULEC (Fig 4F) and HPMEC cells (Fig S4C). Interestingly, Kp43816 induced Thr^495^ phosphorylation even in cells treated with eNOS agonists (Fig S4D). However, the phosphorylation of this site was not observed in cells infected with the *tssB* and *vgrG4* mutants (Fig S4E and Fig 4F). Furthermore, infection with YeVgrG4 also triggered the phosphorylation of Thr^495^ (Fig 4G). Altogether, this evidence establishes that VgrG4 is required to induce the phosphorylation of the eNOS inhibitory site Thr^495^ to blunt eNOS activity.

Collectively, the results demonstrate that *Klebsiella* blunts eNOS activity by inducing the phosphorylation of the inhibitory site Thr^495^ in a T6SS VgrG4-dependent manner and limiting the phosphorylation of the activation site Ser^1177^ in a CPS-dependent manner. Notably, the T6SS VgrG4-dependent phosphorylation of Thr^495^ alone should suffice to halt eNOS activity, thereby leading to the inhibition of vasodilation^39–41^.

### CPS limits eNOS Ser^1177^ phosphorylation by activating the phosphatase PP2Ac in an EGFR-dependent manner

We sought to determine how *Klebsiella* inhibits Ser^1177^ phosphorylation in a CPS-dependent manner. The fact that Kp43816 inhibits the site phosphorylation induced by three different agonists with distinct receptors and modes of action suggests that *Klebsiella* does not target an upstream signalling event related to eNOS activation. We thus considered whether *Klebsiella* may enable a cellular phosphatase to limit the phosphorylation of Ser^1177^. This hypothesis aligns with the observation that drugs inhibiting the phosphorylation of this site activate the phosphatase PP2Ac^41,42^ . Supporting this mechanism, Kp43816 increased the levels of PP2Ac in HULECs (Fig 5A) and HPMECs (Fig S5A). We then investigated whether the CPS is the *Klebsiella* factor responsible for the upregulation of PP2Ac. Immunoblot analysis showed that infection with the *cps* mutant did not raise the levels of PP2Ac (Fig 5A). These findings suggest that CPS-induced upregulation of PP2Ac could be the mechanism exploited by *Klebsiella* to inhibit agonist-induced Ser^1177^ phosphorylation. To confirm this notion, we reduced PP2Ac levels using siRNA and then we assessed the phosphorylation of Ser^1177^ by immunoblotting. *Pp2ac* knockdown efficiency is shown in supplementary figure S5B. In *pp2ac* knockdown cells, Kp43816 did not inhibit ionomycin-induced Ser^1177^ phosphorylation in contrast to cells treated with All Stars control siRNA (Fig 5B). Together, these findings suggest that *Klebsiella* increases the levels of the phosphatase PP2Ac in a CPS-dependent manner to inhibit the phosphorylation of Ser^1177^.

**Figure 5.**
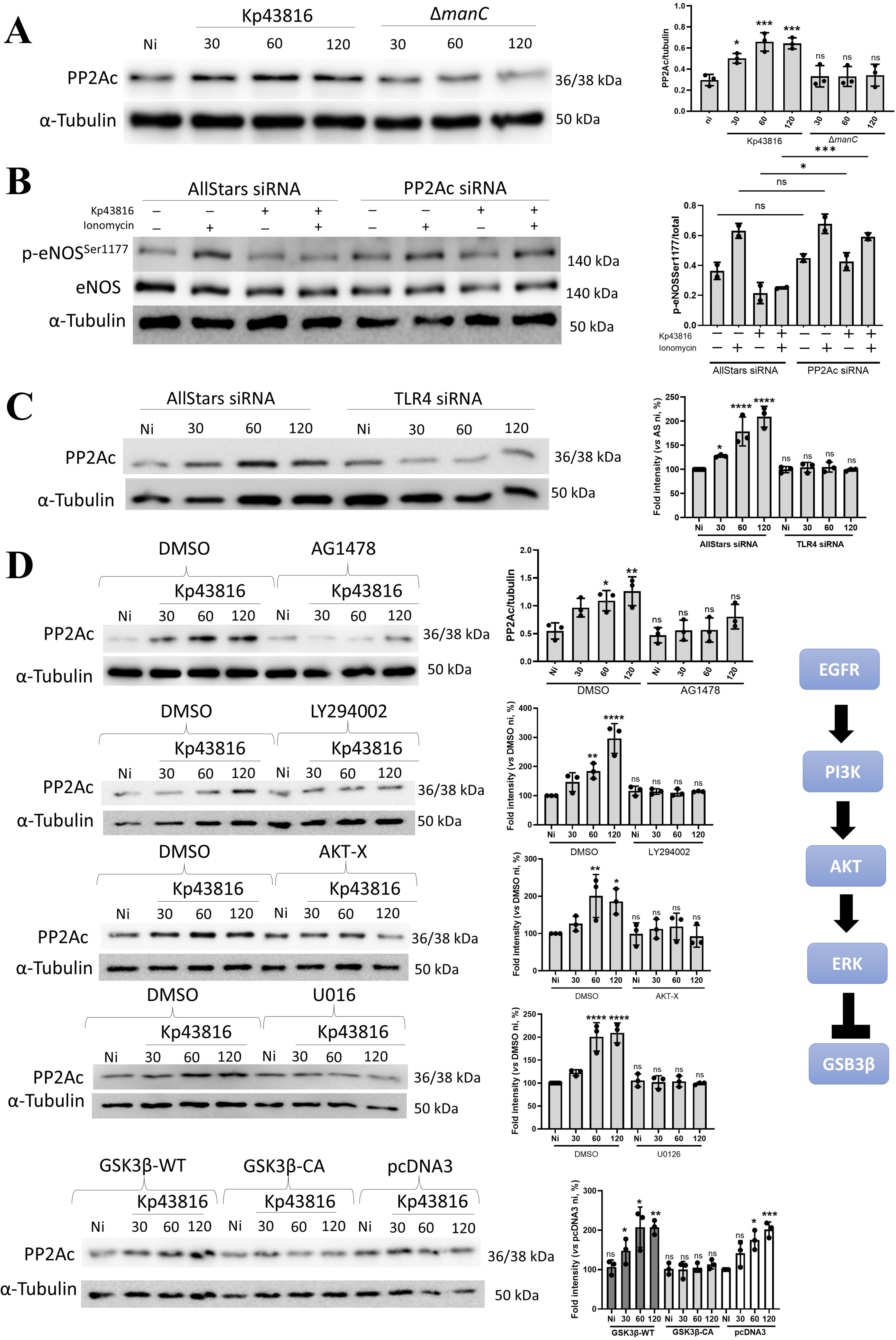
*K. pneumoniae* licences PP2Ac phosphatase to abrogate agonist-induced eNOS Ser^1177^ phosphorylation. (A). Immunoblot analysis of PP2Ac and tubulin levels in HULEC cells infected with Kp43816 and the *cps* mutant (Δ*manC*; 43816-Δ*manC*) for the indicated time points. Densitometry analysis of blots (n = 3) representing the ratio of PP2Ac versus tubulin. (B). Immunoblot analysis of phosphorylated eNOS Ser^1177^ in HULEC cells transfected with All Stars siRNA control, or PP2Ac siRNA. Cells were non-infected, treated for 5 min 1 μM ionomycin, infected for 60 min, or infected and treated with ionomycin. Densitometry analysis of blots (n = 3) representing the ratio of phosphorylated eNOS Ser^1177^ versus eNOS. (C). Immunoblot analysis of PP2Ac and tubulin levels in HULEC cells transfected with All Stars siRNA control, or TLR4 siRNA infected with Kp43816 for the indicated time points. Densitometry analysis of blots (n = 3) representing the ratio of PP2Ac versus tubulin. (D). Immunoblot analysis of PP2Ac and tubulin levels in HULEC cells infected with Kp43816 for the indicated time points. Cells were pretreated with AG1478 (5 μM), LY294002 (20 μM), AKT-X (30 μM), U0126 (10 μM), or DMSO (vehicle solution) for 2 h before infection where indicated. Where indicated, cells were transfected with plasmids expressing GSK3β either wild-type (GSK3β-WT) or with a S9A mutation that renders the protein constitutively active (GSK3β-CA) or with pcDNA3, empty vector. The signalling pathway EGFR-PI3K-AKT-ERK-GSK3β activated in *K. pneumoniae* infection is depicted to represent the several steps inhibited in this figure. Densitometry analysis of blots (n = 3) representing the ratio of PP2Ac versus tubulin. Blots are representative of three independent experiments. In all panels, data were compared against the non-infected control using one-way ANOVA and Tukey’s multiple group comparison correction. The asterisks indicate p,0.05 (*), <0.01 (**). <0.001 (***), or <0.0001 (****). ns, not significant.

To decipher how CPS increases the levels of PP2Ac, we reasoned that the CPS should activate a signalling pathway controlling the levels of the phosphatase. Previously, we demonstrated that CPS activates TLR4 to activate an EGFR-phosphatidylinositol 3-OH kinase (PI3K)-AKT-PAK4-ERK-GSK3β signalling pathway to upregulate the levels of the CYLD deubiquitinase in human lung cells^43^. Therefore, we asked whether this pathway also controls the levels of PP2Ac in human endothelial cells. We first tested whether Kp43816 activates this pathway in human endothelial cells. In *tlr4* knockdown cells, Kp43816 did not phosphorylate EGFR (Fig S5C). *tlr4* knockdown efficiency was higher than 70 % (Fig S5D). Immunoblot analysis confirmed that Kp43816 activates the phosphorylation of the nodes of the pathway AKT, PAK4, and ERK (Fig S5E). GSK3β is a constitutively active kinase under resting conditions but becomes inactivated by phosphorylation of serine 9 by AKT^44,45^. As expected, Kp43814 triggered the phosphorylation of GSK3β (Fig S5E). In *tlr4* knockdown cells, Kp43816 did not increase the levels of PP2Ac (Fig 5C). We utilized pharmacologic inhibitors of EGFR (AG1478), PI3K (LY294002), AKT (AKT-X) or MEK (U0126) to block the pathway at four different levels. Results shown in Figure 5D revealed that Kp43186-induced PP2Ac expression was reduced in cells pretreated with these inhibitors. To examine the role of inhibitory phosphorylation of GSK3β on Kp43816-controlled upregulation of PP2Ac, we studied the effect of forced expression of GSK3β. Vectors encoding HA tag wild-type GSK3β (GSK3β-WT) or a constitutive active mutant (GSK3β-CA) in which the serine 9 residue was changed to alanine were transiently transfected to HULEC cells. As control, cells were transfected with the empty vector pcDNA3. Kp43816 did not increase the expression of PP2Ac in cells transfected with GSK3β-CA vector (Fig 5D).

In summary, the evidence presented supports the notion that *Klebsiella* activates an TLR4-EGFR-PI3K-AKT-PAK4-ERK-GSK3β signalling pathway in a CPS-dependent manner to increase the levels of the phosphatase PP2Ac to inhibit the phosphorylation of eNOS Ser^1177^.

### VgrG4 induces eNOS Thr^495^ phosphorylation by activating the kinase PKCβ in a NLRX1-ROS-dependent manner

Structure-function studies of VgrG4 do not indicate that VgrG4 has any kinase activity^32^. To explain VgrG4-stimulated Thr^495^ phosphorylation we speculated that VgrG4 licenses the activity of a cellular kinase implicated in the phosphorylation of Thr^495^. Different kinases of the PKC family phosphorylate Thr^495^ in conditions such as hypoglycaemia and atherosclerosis^39,41,46–48^. We therefore asked whether *Klebsiella* activates PKC. Infection of HULEC and HPMEC cells with Kp43816 triggered the phosphorylation of the active site common to all isoforms of the PKC family (Fig 6A and Fig S6A). To connect the activation of PKC with *Klebsiella*-induced phosphorylation of the Thr^495^ eNOS inhibitory site, infections were done in the presence of the PKC inhibitor Ro31-8220. In these conditions, we did not detect the phosphorylation of Thr^495^ (Fig S6B). To identify the PKC isoform mediating Thr^495^ phosphorylation we carried out a siRNA-based screen probing Thr^495^ phosphorylation upon infection by immunoblotting. Of the 3 PKC isoforms tested, PKCα, β, and ε, Kp43816-induced Thr^495^ phosphorylation was only abrogated in cells in which PKCβ was knocked down (Fig 6B and Fig S6C). Control experiments confirmed the reduction of the *pkc* transcripts in HULEC cells (Fig S6D). Thr^495^ phosphorylation was also not observed in infected cells treated with the PKCβ inhibitor LY333-531^49^ (Fig S6E). YeVgrG4 did not trigger Thr^495^ phosphorylation in PKCβ knockdown cells (Fig 6C). Altogether, these results support the notion that VgrG4 triggers the activation of PKCβ to phosphorylate Thr^495^, thereby abrogating eNOS activity.

**Figure 6.**
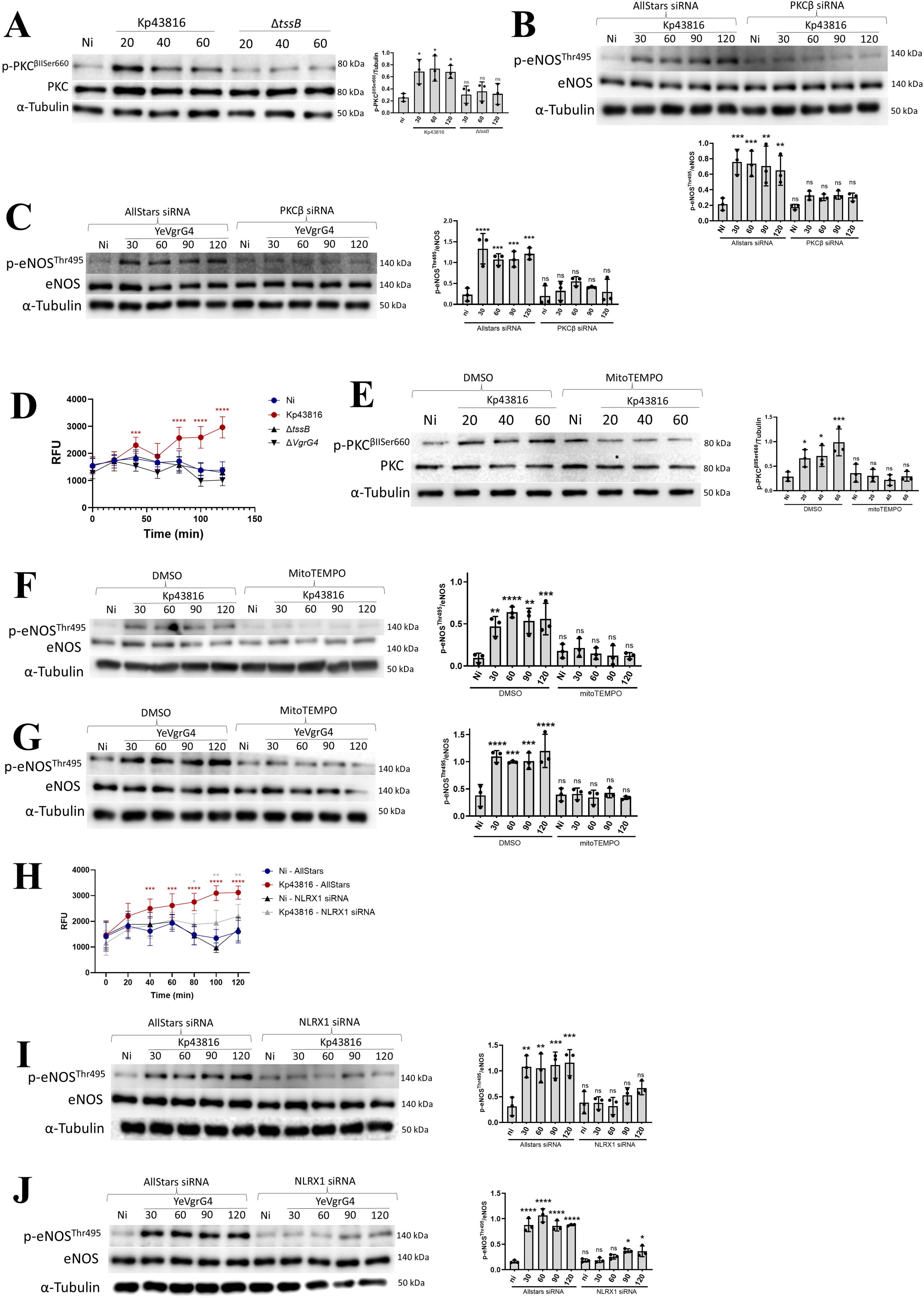
*K. pneumoniae* licences PKCβ kinase to phosphorylate eNOS Thr^495^. (A). Immunoblot analysis of phosphorylated PKC and tubulin levels in HULEC cells infected with Kp43816 and T6SS mutant *tssB* (Δ*tssB*; 43816-Δ*tssB*) for the indicated time points. Densitometry analysis of blots (n = 3) representing the ratio of phosphorylated PKC versus tubulin. (B). Immunoblot analysis of phosphorylated eNOS Thr^495^, eNOS and tubulin levels in HULEC cells transfected with All Stars siRNA control, or PKCβ siRNA infected with Kp43816 for the indicated time points. Densitometry analysis of blots (n = 3) representing the ratio of phosphorylated eNOS Thr^495^ versus eNOS. (C). Immunoblot analysis of phosphorylated eNOS Thr^495^, eNOS and tubulin levels in HULEC cells transfected with All Stars siRNA control, or PKCβ siRNA infected with YeVgrG4 for the indicated time points. Densitometry analysis of blots (n = 3) representing the ratio of phosphorylated eNOS Thr^495^ versus eNOS. (D). mtROS was quantified as relative fluorescence units (RFU) in HULEC cells treated with MitoSOX (2 µM, 45 min before infection) and then infected with Kp43816, and the T6SS *tssB* (Δ*tssB*; 43816-Δ*tssB*) and *vgrG4* (Δ*vgrG4*; Kp43816-Δ*vgrG4*) mutants for the indicated time points. (E). Immunoblot analysis of phosphorylated PKC, PKC and tubulin levels in HULEC cells treated with vehicle solution or MitoTEMPO and infected with Kp43816 for the indicated time points. Densitometry analysis of blots (n = 3) representing the ratio of phosphorylated PKC versus tubulin. (F). Immunoblot analysis of phosphorylated eNOS Thr^495^, eNOS and tubulin levels in HULEC cells treated with vehicle solution or MitoTEMPO, and infected with Kp43816 for the indicated time points. Densitometry analysis of blots (n = 3) representing the ratio of phosphorylated eNOS Thr^495^ versus eNOS. (G). Immunoblot analysis of phosphorylated eNOS Thr^495^, eNOS and tubulin levels in HULEC cells treated with vehicle solution or MitoTEMPO, and infected with YeVgrG4 for the indicated time points. Densitometry analysis of blots (n = 3) representing the ratio of phosphorylated eNOS Thr^495^ versus eNOS. (H). mtROS was quantified as RFU in HULEC cells transfected with All Stars siRNA control, or NLRX1 siRNA treated with MitoSOX (2 µM, 45 min before infection) and then infected with Kp43816 for the indicated time points. (I). Immunoblot analysis of phosphorylated eNOS Thr^495^, eNOS and tubulin levels in HULEC cells transfected with All Stars siRNA control, or NLRX1 siRNA infected with Kp43816 for the indicated time points. Densitometry analysis of blots (n = 3) representing the ratio of phosphorylated eNOS Thr^495^ versus eNOS. (J). Immunoblot analysis of phosphorylated eNOS Thr^495^, eNOS and tubulin levels in HULEC cells transfected with All Stars siRNA control, or NLRX1 siRNA infected with YeVgrG4 for the indicated time points. Densitometry analysis of blots (n = 3) representing the ratio of phosphorylated eNOS Thr^495^ versus eNOS. Blots are representative of three independent experiments. In panels D and H, data is shown as mean ± SD of three independent experiments. In all panels for densitometry, data were compared against the non-infected control using one-way ANOVA and Tukey’s multiple group comparison correction. In panels D and H, data were compared against non-infected control using two-way ANOVA and Dunnet’s multiple group comparison correction. The asterisks indicate p<0.05 (*), <0.01 (**). <0.001 (***), or <0.0001 (****). ns, not significant.

Different signals have been reported to activate PKCβ, with ROS being one of them^50^. Interestingly, VgrG4 induces mitochondrial ROS (mtROS) in human epithelial cells^26^, suggesting that VgrG4-induced mtROS may activate PKCβ in endothelial cells. We first ascertained whether VgrG4 induces mtROS in HULEC cells by assessing the oxidation of the mitochondria localized fluorescent dye MitoSOX. Kp43816 induced mtROS in HULEC cells in a T6SS VgrG4-dependent manner because, unlike the wild-type strain, neither the *tssB* nor the *vgrG4* mutants increased the levels of mtROS over those of the non-infected cells (Fig 6D). To substantiate that VgrG4-induced mtROS triggers the activation of PKCβ to phosphorylate Thr^495^, infections were done in the presence of the mtROS inhibitor MitoTEMPO. Scavenging mtROS with MitoTEMPO prevented Kp43816-induced activation of PKC (Fig 6E). Furthermore, Kp43816 (Fig 6F) and YeVgrG4 (Fig 6G) did not induce Thr^495^ phosphorylation in MitoTEMPO-treated cells, demonstrating that mtROS mediates the activation of PKCβ.

We have recently demonstrated that VgrG4 induction of mtROS is dependent on the activation of the mitochondria located innate receptor NLRX1 in human lung epithelial cells^26^. Quantification of MitoSOX oxidation corroborated that Kp43816 infection did not increase the levels of mtROS in *nlrx1* knockdown cells using siRNA (Fig 6H), establishing that NLRX1 activation also controls *Klebsiella*-induced mtROS in human endothelial cells. *nlrx1* knockdown efficiency is shown in supplementary figure S6F. In *nlrx1* knockdown cells, Kp43816 (Fig 6I) and YeVgrG4 (Fig 6J) did not trigger the phosphorylation of Thr^495^, demonstrating that Thr^495^ phosphorylation by *Klebsiella* VgrG4 is NLXR1-dependent.

Altogether, this evidence supports the notion that VgrG4-induced NLRX1-dependent mtROS activates PKCβ to subsequently trigger the phosphorylation of the eNOS inhibitory site Thr^495^ to blunt eNOS activity.

### Inhibition of mtROS and PKCβ restores ACh-induced vasodilation in *K. pneumoniae*-infected vessels

Based on our cell culture studies and the importance of Thr^495^ phosphorylation in the inhibition of eNOS activity^38,39^, we next aimed to determine whether inhibiting mtROS and PKCβ would restore ACh-induced vasodilation in infected vessels. Initially, we examined the impact of mtROS inhibition. MitoTEMPO did not affect ACh-induced vasodilation in controls vessels (Fig 7A and Fig S7A). However, treatment with MitoTEMPO restored ACh-induced vasodilation in Kp43816-infected vessels to levels comparable to those observed in control vessels (Fig 7A and Fig S7B). This restoration of ACh-induced vasodilation involved both the NO and EDH pathways, as evidenced by experiments delineating these pathways using L-NAME, TRAM-34, and apamin (Fig 7A and Fig S7C and Fig S7D). These findings demonstrate that inhibition of VgrG4-induced mtROS restores the NO and EDH pathways affected by *K. pneumoniae* infection.

**Figure 7.**
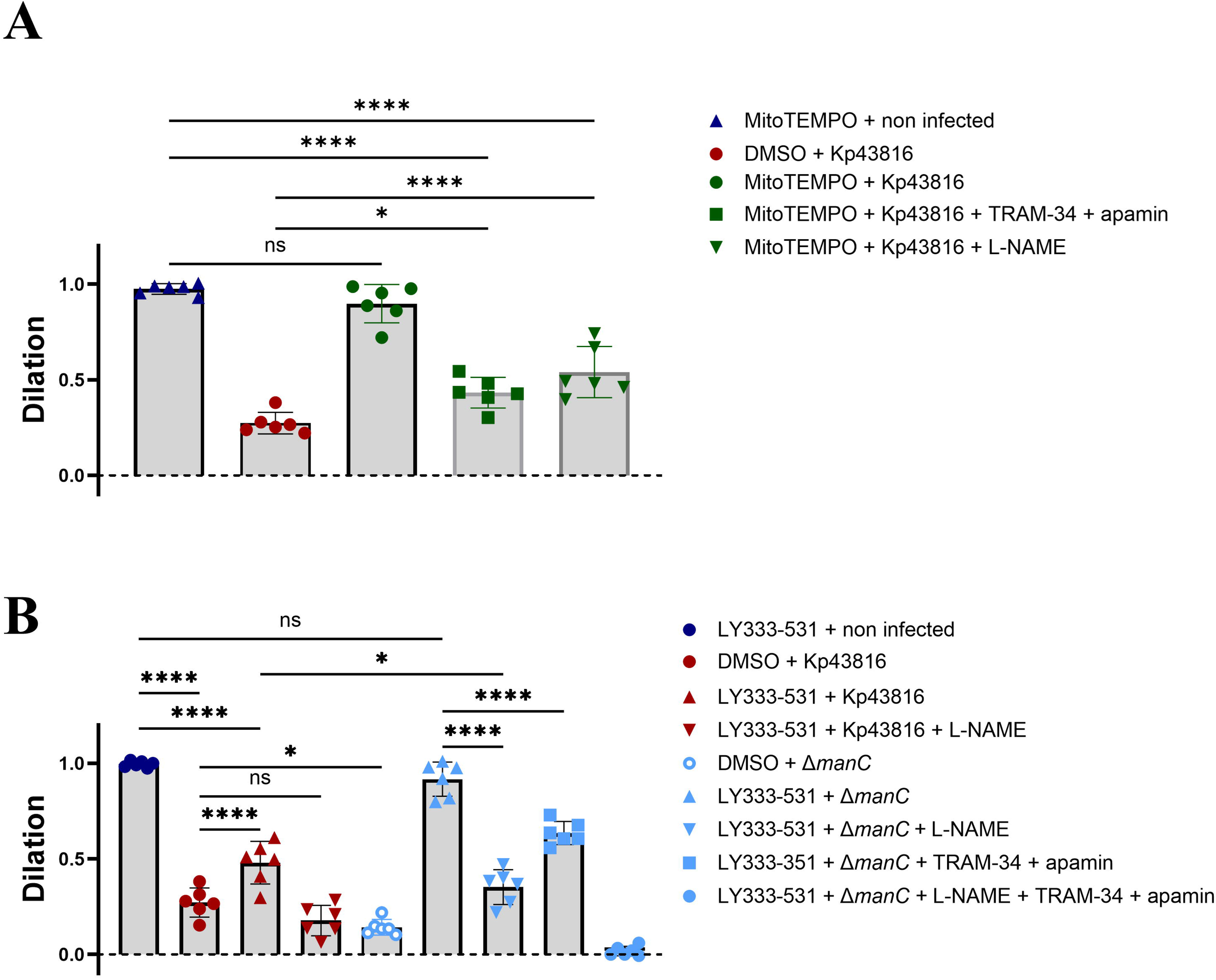
Inhibition of mtROS and PKCβ restores ACh-induced vasodilation in vessels infected with *K. pneumoniae*. (A). ACh-induced peak dilation in vehicle solution, and MitoTEMPO treated vessels infected with Kp43816. When indicated, infected vessels were also treated with the NO pathway inhibitor L-NAME, and the EDH pathway inhibitors TRAM-34 and apamin. (B). ACh-induced peak dilation in vehicle solution, and PKCβ inhibitor LY333-531 treated vessels infected with Kp43816, and the *cps* mutant (Δ*manC*; 43816-Δ*manC*). When indicated, infected vessels were also treated with the NO pathway inhibitor L-NAME, and the EDH pathway inhibitors TRAM-34 and apamin. In A and B, ACh was used at 1 μM, and peak dilation is shown as mean ± SD from six vessels from three rats per group and analysed with two-way ANOVA and Dunnet’s multiple group comparison correction. The asterisks indicate p < 0.05 (*), or <0.0001 (****). ns, not significant.

We next tested the effect of inhibiting PKCβ in infected vessels. Control experiments showed that the PKCβ inhibitor LY333-531 did not affect ACh-induced vasodilation in controls vessels (Fig 7B and Fig S7E). In contrast, we observed a significant increase in vasodilation in infected vessels treated with LY333-531 (Fig 7B and Fig S7F). This ACh-induced vasodilation was mediated by the NO signalling pathway because L-NAME abrogated ACh-triggered vasodilation in infected vessels treated with LY333-531 (Fig 7B and Fig S7G). The fact that PKCβ inhibition did not restore fully the vasodilation of infected vessels to that of control vessels led us to consider whether this could be mediated by the CPS effect on Ser^1177^. If this is the case, then full vasodilation should be observed in vessels infected with the *cps* mutant and treated with LY333-531. Indeed, this was the case (Fig 7B and Fig S7H). Inhibition with L-NAME, and TRAM-34 and apamin significantly reduced ACh-induced vasodilation (Fig 7B and Fig S7I and Fig S7J). As expected, combination of L-NAME with TRAM-34 and apamin abolished ACh-triggered vasodilation in infected vessels with the *cps* mutant treated with LY333531 (Fig 7B and Fig S7K).

Altogether, these findings illustrate the physiological significance of inhibiting *Klebsiella*-controlled actions on eNOS phosphorylation at Ser^1177^ and Thr^495^, resulting in the restoration of vasodilation.

## DISCUSSION

Here, we establish a research platform, an ex vivo blood vessel model coupled with challenging human primary endothelial cells, to dissect how pathogens affect vascular physiology. We demonstrate that *K. pneumoniae* inhibits vasodilation by blunting the NO-dependent pathway and attenuating the EDH pathway (Fig S8). Mechanistically, *Klebsiella* exploits the trans-kingdom T6SS effector VgrG4 to trigger the phosphorylation of the eNOS inhibitory site Thr^495^ by licensing the kinase PKCβ in an NLRX1-mtROS-dependent manner (Fig S8). Additionally, *Klebsiella* inhibits the phosphorylation of the eNOS activation site Ser^1177^ by licencing the phosphatase PP2Ac upon activation of an EGFR-dependent pathway in a CPS-dependent manner (Fig S8). Our results also indicate that VgrG4-induced mtROS attenuates the EDH pathway. Collectively, our findings unveil hitherto unknown anti-host functions of the T6SS and the CPS resulting in endothelial dysfunction. Altogether, this work illustrates a new axis exploited by a human pathogen to govern vascular physiology.

The research platform described in this work allowed us to dissect an angle of the host-*Klebsiella* interaction never investigated before despite the global clinical relevance of *Klebsiella*-caused BSI infections. In fact, this is a poorly understood niche in the host-pathogen arms race for most bacterial pathogens causing BSI. To date the research in this area has mostly focused on investigating the interaction of pathogens with endothelial cells from the vein of the umbilical cord or the retina. This research has generated valuable observations, yet with limited insights into how the findings translate into vascular physiology. In contrast, our ex vivo blood vessel model enables the study and quantification of vascular physiology upon infection. Moreover, here we have challenged human relevant endothelial cells to decipher the molecular signalling pathways targeted by *Klebsiella*. The fact that the findings using these cells were translatable to effects on vascular physiology in our ex vivo model underscores the robustness of our research framework.

Inhibition of vasodilation impairs the perfusion of infected tissue reducing the recruitment of immune cells into the surrounding tissue and, therefore, it is considered a pathophysiological feature of infections particularly in the case of viral infections. In sharp contrast to viral and bacterial infections that induce endothelial dysfunction by damaging the endothelial barrier^51,52^, we have demonstrated that *K. pneumoniae* causes endothelial dysfunction by reducing the availability of NO without significantly disturbing the endothelial layer. The importance of NO in the outcome of *Klebsiella* infection is marked by the fact that inhibition of NO production using L-NAME results in increased mortality^53^. The key molecular mechanism behind *Klebsiella*-induced inhibition of vasodilation involves altering eNOS posttranslational phosphorylation. While Thr^495^ and Ser^1177^ are well characterized targets of inhibitors and agonists of vascular vasodilation, until now, there has been no evidence to suggest that pathogens commonly exploit these targets. As far as we know, *K. pneumoniae* appears to be the first pathogen to target both regulatory sites. Mechanistically, *Klebsiella* licences PKCβ kinase and the PP2Ac phosphatase to exert its effects on eNOS phosphorylation. Both enzymes govern the homeostatic regulation of eNOS activation^54^, illustrating a strategy radically different to those of other pathogens such as *Salmonella, Legionella, Escherichia* and *Shigella*, which deploy bacterial proteins to directly target host proteins. Previous work from our laboratory demonstrated that *Klebsiella* also exploits the deubiquitinase, CYLD, the phosphatase MKP-1, the deSUMOylase SENP2, and the deneddylase CSN5, to affect host protein post translational modifications^43,55,56^. Cells activate CYLD and MKP-1 to return to homeostasis after inflammation to protect the host from an overwhelming inflammatory response^57,58^, whereas SENP2 and CSN5 catalyse the deconjugation of SUMO and NEDD8, respectively, from proteins to control signalling pathways modulated by SUMOylation and NEDDylation^59,60^. Therefore, a discernible theme emerges whereby a key aspect of *K. pneumoniae* infection biology involves licencing host proteins responsible for restoring cellular homeostasis, thereby manipulating protein post translational modifications to alter cell biology.

Another novel finding of this work is that the T6SS controls vascular physiology, expanding substantially the capabilities of this secretion system. This new anti-host function of *Klebsiella* T6SS adds to its role blocking NF-κB signalling through mitochondrial alteration^26^, governing the landscape of lung myeloid cells and their interactions with *Klebsiella* to promote lung infection^27^, facilitating gut colonization^61^, and its antimicrobial role against bacteria and fungi^32^.The trans-kingdom effector VgrG4 mediates these anti-host activities although we anticipate there are other *Klebsiella* T6SS effectors with anti-host function considering the significant diversity of effectors found in *K. pneumoniae* T6SS^32,62^. Future work is warranted to define the portfolio of T6SS antimicrobial toxins and those with anti-host function.

VgrG4-exerted inhibition of vasodilation by targeting eNOS Thr^495^ is dependent on NLRX1-controlled mtROS to activate PKCβ. *Klebsiella* also exploits NLRX1-induced mtROS to inhibit the activation of NF-κB signalling with a concomitant reduction in inflammation^26^. The role of NLRX1 on infection remains controversial. Research has focused on assessing its role in inflammation, acting as a positive or negative regulator depending on the infection context^63–66^. Likewise, there is conflicting data on the role of NLRX1 on cell death^67–70^. In contrast, our research places NLRX1 unequivocally at the fulcrum of the strategies deployed by *Klebsiella* to counteract the host. Furthermore, our work sheds light on the role of NLRX1 in vascular physiology, laying the groundwork for future investigations into its contribution to diseases characterized by endothelial dysfunction resulting from deficiencies in NO bioavailability.

Whereas inhibition of mtROS-induced activation of PKCβ is sufficient to restore the NO pathway in infected vessels, only when mtROS is inhibited is the EDH pathway is restored. Together, these findings indicate that mtROS is the main signal governing VgrG4-inhibition of vasodilation affecting the NO and the EDH pathways. How VgrG4-induced mtROS affects the EDH pathway remains an open question. Our results ruled out that mtROS may affect the levels of intracellular Ca^2+^ required for the activation of SKCa and IKCa channels. We speculate, however, that mtROS may influence channel activity by reducing their open probability in response to rises in intracellular Ca^2+^, thereby reducing endothelial hyperpolarization. It is also possible that mtROS may negatively impact the transmission of this signal through myoendothelial gap junctions. Future studies will dissect the molecular mechanism of mtROS-dependent attenuation of the EDH pathway.

Although VgrG4 is necessary and sufficient to inhibit vasodilation, we have shown that the CPS also contributes to this phenotype by preventing the phosphorylation of the active site of eNOS in an EGFR-dependent manner, highlighting the importance of inhibiting eNOS activity for *Klebsiella* infection biology. The work of the laboratory and others have established that *Klebsiella* CPS is a bona fide immune evasin^25^. CPS contributes to *Klebsiella* stealth strategy by limiting the recognition of the pathogen by innate receptors^71^. In addition, it activates EGFR-dependent signalling to dampen NF-κB activation, thus attenuating inflammation through the inhibition of K-63 ubiquitination of TRAF6 by the ubiquitinase CYLD^43,55^. *Klebsiella* also restricts two other post translational modifications, SUMOylation and NEDDylation, through activation of this EGFR-dependent pathway^56^. Altogether, this evidence supports the notion that *Klebsiella* controls the post translational modifications of host proteins via this EGFR-governed pathway.

Host-directed therapeutics are gaining momentum as a viable approach to treat infectious diseases^72^. It is interesting to note that *Klebsiella*-induced endothelial dysfunction resembles that seen in cardiovascular diseases like hypertension and atherosclerosis, opening the opportunity to explore whether some of the therapeutics for these conditions could also be effective in treating *K. pneumoniae* infections, either alone or as a synergistic complement to antibiotic treatment. Future studies shall confirm whether this is the case.

## METHODS

### Bacterial strains and growth conditions

Strains and bacterial mutants are shown in Table S1. *K. pneumoniae* strains and mutants were cultured over 16-18 hours in 5 ml LB and refreshed the next day by the addition of 0.5 ml of the overnight culture into 4.5 ml of fresh medium. Bacteria were harvested at mid exponential phase (2500 × g, 20 min) and adjusted to an optical density of 1.0 at 600 nm (OD_600_) in PBS (5 × 10^8^ CFU/ml). *Y. enterocolitica* strains were grown in 5 ml LB overnight and refreshed in tryptic soy broth (TSB) media containing 20 mM MgCl_2_ and 20 mM sodium oxalate for 3.5 h hours at 21°C followed by 30 minutes at 37°C to activate *Yersinia* T3SS. Bacteria were recovered by centrifugation and adjusted to an OD_600_ of 1.0 in PBS (5 × 10^8^ CFU/ml).

### Ex vivo blood vessel and infection

Male Sprague Dawley rats between the age of 6-12 weeks were culled using a Schedule 1 method and their mesenteric arcade was isolated and placed into ice cold PSS [118 mM NaCl, 4.7 mM KCl, 20 mM HEPES, 5 mM NaHCO_3_, 1.2 mM MgCl_2_, 1.2 mM KH_2_PO_4_, 2 mM CaCl_2_ and 5 mM glucose]. The mesenteric arcade was pinned onto a sylgard coated dish using insect pins. Using vannas scissors and microdissection forceps, 2^nd^ and 3^rd^ order mesenteric arteries were separated from the adipose tissue and placed into low calcium PSS [118 mM NaCl, 4.7 mM KCl, 20 mM HEPES, 5 mM NaHCO_3_, 1.2 mM MgCl_2_, 1.2 mM KH_2_PO_4_, 100 μM CaCl_2_ and 5 mM glucose], and stored at 4°C. Vessels over 250 μm, under no pressure, were discarded. Each vessel was cannulated using pipettes pulled with a Stutter instrument P-97, using thin-walled borosilicate glass capillaries with an outer diameter of 1.5 mm. Vessels were tied down on the cannula using two nylon size 8/0 sutures half knotted on each end within a DMT Culture Myograph 204 chamber. The myograph chamber contained PSS and the vessel was perfused with PSS. The chamber and the PSS were kept at 37°C.

After cannulation, vessels were allowed to equilibrate for 20 minutes at an intraluminal pressure of 70 mmHg. Subsequently, 10 μM PE was added into the chamber to induce a sustained constriction, which was typically achieved within 2-3 minutes. As indicated in the traces, ACh was added to the chamber in a step-wise (concentrations from 1 nM to 100 μM). When the vessels were infected, bacteria were flowed into the vessel and then the flow was stopped. Vessels were infected for 1-hour with *K. pneumoniae* strains or 30 minutes with *Y. enterocolitica* prior to addition of PE and ACh. The outer diameter of the vessel was imaged and recorded using an infrared microscope (DMT). The traces were reproduced using the DMT Myoview 5 software.

NO pathway-dependent dilation was inhibited with 100 μM L-NAME whereas EDH pathway-controlled dilation was inhibited with 1 μM TRAM-34 and 100 nM apamin. Inhibitors were added to the chamber 30 min before infection and maintained during the duration of the experiment. To ascertain smooth muscle cell reactivity to NO, 5 min after addition of 10 μM PE, 100 μM SNP was added to the chamber. In the experiments testing the effect of inhibiting PKCβ and mtROS, vessels were incubated with 100 nM of LY333-531 and 10 μM MitoTEMPO, respectively 1-hour before infection and maintained during the duration of the experiment.

All myography experiments were performed probing two vessels per rat. Vessels from three rats were tested unless stated otherwise.

### Calcium signalling in endothelial cells

To quantify intracellular Ca^2+^ signals in cultured endothelial cells in response to agonists and infection, we employed Fura2-QBT assays (Molecular Devices). HULEC cells were seeded in 96 well black walled, clear bottom plates, at 20,000 cells per well, 24 hours before the experiment. To load cells with the dye Fura2QBT, wells were washed extensively with HBSS to remove cell media and loaded with 200 μl Fura2QBT in HBSS. Cells were incubated for 1 h at 37°C before addition of drugs. Drugs were prepared in a 96 well V-bottom master plate in HBSS at 4x the final concentration. The plates were placed into a temperature controlled Flexstation 3 system at 37°C, which included custom manufactured tips (Molecular Devices) that were used to pipette 50 μl of the drugs at predetermined settings (as per manufacturer guidance). To assess the effect of infection, cells were infected at a multiplicity of infection 100 bacteria per cell in HBSS after loading the cells with Fura2QBT. After 2 h, the plates were placed in the Flexstation 3 system for the addition of the drugs as previously described.

Intracellular Ca^2+^ levels were assessed for 30 seconds before and for 5 minutes following the addition of the drugs. Excitation was performed at 340/380 nm, with emission measurements recorded at 510 nm. All experiments were performed with six technical replicates, in three independent occasions.

### Ex vivo measurement of Ca^2+^ responses by the endothelium

Small rat mesenteric resistance arteries (2^nd^ and 3^rd^ order) were cut open longitudinally and pinned out using tungsten wire, exposing the endothelium using microdissection scissors. Vessels were loaded with Cal520-AM (5 µM with 0.02% Pluronic F-127 in PSS at 37°C for 30 min). After incubation with the dye, the chamber was washed with PSS to remove excess dye. A custom-made chamber was placed on an upright fluorescence microscope and the endothelium was imaged with a 16x (0.8 NA) water immersion objective lens at 10Hz (488nm excitation wavelength). All images were captured using a Photometrics Evolve 13 EMCCD camera (1024 x 1024) camera, and the data recorded with μManager v2 software.

Endothelium Ca^2+^ responses were measured before and after addition of ACh in step-wise increments (3 nM, 100 nM and 3 μM) in control vessels. After completing the recording with the last concentration of ACh, the vessel was thoroughly washed, and then mock-infected or infected with 10^6^ CFU/ml Kp43816 for 1 h before the same concentrations of ACh were added. At the end of the experiment, 4-DAMP (M3 muscarinic receptor antagonist) was added to demonstrate that the ACh -induced Ca^2+^ signalling was dependent on the activation of M3 muscarinic receptors. All results for Ca^2+^ signalling experiments for mock-infected and infected vessels were normalised to their respective vessel control values. Data was plotted using a published algorithm^73^.

In these experiments, a total of six vessels from six different rats were probed.

### Construction of *vgrG4* mutant

Primers for mutant construction (Table S2) were designed based on the whole-genome sequence of *K. pneumoniae* ATCC43816 (GenBank accession no. CP009208.1). The primer pairs VgrG4 UP and VgrG4 DOWN were used in separate PCR reactions using Q5 DNA Polymerase (New England Biolabs) to amplify a 730 bp and a 1027 bp fragments flanking the *vgrG4* gene. BamHI restriction sites internal to these flanking regions were incorporated at the end of each amplicon. Purified *vgrG4* UP and DOWN fragments were then polymerised and amplified as a single PCR amplicon using the primers vgrG4_UPFWD and vgrG4_DWNRVS. The 1.7 kb PCR amplicon was gel-purified and cloned by Homologous Alignment Cloning^74^ into EcoRI-digested Antarctic Phosphatase (New England Biolabs)-treated and gel-purified pFOK suicide vector^75^ to generate pFOKVgrG4. Plasmid was transformed by heat shock into *E. coli* JKe201 and transformants were selected in on LB agar plates containing 100 μM DAP (required for growth of JKe201) and kanamycin (50 μg/ml). pFOKVgrG4 was mobilised to *K. pneumoniae* ATCC43816 by conjugation as previously described^75^, and co-integrants were selected on LB plates containing kanamycin (50 μg/ml). At least three co-integrant clones were combined and grown for 4 h at 37 °C in 2 ml of LB. Bacteria were then streaked on freshly prepared LB-no salt agar plates containing 20% sucrose and 0.5 μg/ml anhydrous tetracycline. Plates were incubated at 28 °C protected from light for at least 24 h. Candidate mutant clones were confirmed by PCR using VgrG4 SCREEN primer pair. The confirmed mutant was named Kp43816-Δ*vgrG4*.

### Assessment of bacterial growth

To test the effect of inhibitors on the growth of *Klebsiella*, 5 μl from an overnight culture in a 96-well plate were diluted in 250 μl of LB or M9 minimal medium (5× M9 minimal salts [Sigma-Aldrich] supplemented with 2% glucose, 3 mM thiamine, 2 mM MgSO_4_) and incubated at 37°C with continuous, normal shaking in a Bioscreen C Automated Microbial Growth Analyzer (MTX Lab Systems, Vienna, VA, USA). OD_600_ was measured and recorded every 20 min. Results, reported in Fig S9, demonstrate that the inhibitors probed did not affect the growth of *Klebsiella*.

### Cell culture

Immortalised human lung microvascular endothelial cells HULEC5a (ATCC) were grown in glutamine free MCDB131 media (ThermoFisher Scientific) supplemented with 10 mM glutamine, 10mM HEPES (Sigma), 1µg/ml hydrocortisone, 10 ng/ml human epidermal growth factor and 10% FBS. Unless otherwise indicated, cells were seeded at 125,000 per well in a 12 well plate without antibiotic, 24 h before infection. Cells were used between passages 3-10.

Primary human lung microvascular endothelial cells (HPMEC) (Innoprot) were grown in Endothelial cell medium (ECM), supplemented with 10% FBS and endothelial cell growth supplement (Imoprot).

HULEC and HPMEC were infected at a multiplicity of infection of 100 bacteria per cell.

Cells were routinely tested for *Mycoplasma* contamination.

### Transfection and knockdown efficiency

For transfections, cells were seeded at 60,000 cells per well in a 12 well plate, 48 h prior to infection. siRNA transfection was performed using Lipofectamine 2000 (Invitrogen) as per the manufacturer’s instructions (3 μl per 100 μl Optimem), with 20 nM siRNA or negative control siRNA (AllStars - Qiagen), to a final volume of 1ml per well. Knockdown efficiency was confirmed by immunoblotting or by qPCR analysis of duplicate samples from three independent transfections by normalising to the glyceraldehyde 3-phosphate dehydrogenase (hGAPDH) gene and comparing gene expression in the knockdown sample with the AllStars siRNA control. Primers used are listed in Table S2.

### Immunoblot

HULEC and HPMEC cells were seeded in 12 well plates at a density of 125,000 and 80,000 cells respectively, 24 h prior to infection. Infections for western blot analysis were performed with *K. pneumoniae* or *Y. enterocolitica* for various time points as indicated in the figure legends. Where inhibitors or siRNA were used, the details are provided in the figure legends. Cells were then washed in 1 mL of ice-cold PBS, and were lysed in 80 μL of 2 x SDS sample buffer (1x SDS Sample Buffer, 62.5 mM Tris-HCl pH 6.8, 2% w/v SDS, 10% glycerol, 50 mM DTT, 0.01% w/v bromophenol blue). Cell lysates were sonicated for 10 seconds at 10% amplitude with a Branson 450 digital sonicator and were then stored at -20°C until use. Samples were boiled at 95°C for 5 minutes and centrifuged at 12,000 × g for 1 min. 25% (20µl) of the lysates were resolved by standard 10% SDS-PAGE and electroblotted onto nitrocellulose membranes using semi dry Trans-Blot (Bio-Rad). Membranes were blocked with 3% BSA (w/v) in TBST for 1 hour before being incubated overnight at 4°C with the primary antibodies. Immunoreactive bands were visualized by incubation with horseradish peroxidase-conjugated goat anti-rabbit immunoglobulins (1:5000, BioRad 170-6515), or goat anti-mouse immunoglobulins (1:5000, BioRad 170-6516). Bands were detected using chemiluminescence reagents in a G:BOX Chemi XRQ chemiluminescence imager (Syngene).

To detect multiple proteins, membranes were reprobed after stripping of previously used antibodies using a pH 2.2 glycine-HCl/SDS buffer. To ensure that equal amounts of proteins were loaded, blots were reprobed with mouse anti-human tubulin (1:3000, Sigma T5168).

### Densitometry analysis

Images of the blots were exported from the GeneSys software (Syngene) and analysed in Image studio lite (Licor) to obtain signal intensity of the bands. When analysing phosphorylated protein, densitometric analysis was performed by dividing the phosphorylated protein signal intensity over the total protein signal intensity (phospho/total). In cases where total protein was observed, the total protein signal intensity was divided by the signal intensity of the loading control (tubulin).

### Mitochondria ROS detection

HULEC cells were seeded in a 96 well plate (black, clear bottom) at 15,000 cells per well and incubated with 2 µM MitoSOX red for 45 minutes in antibiotic free Hanks’ Balanced Salt Solution (HBSS). Cells were then gently washed three times with prewarmed PBS and infected in fresh antibiotic free HBSS with *K. pneumoniae* as previously described. Fluorescence was measured over 2 hours (readings taken every 20 minutes) at Excitation/Emission 544/590 in a POLARStar Omega BMG LabTech plate reader. All experiments were performed with six technical replicates across three independent occasions.

### Histology

Rat 2^nd^ and 3^rd^ order mesenteric arteries were isolated from the mesenteric arcade, cannulated and any remnants of blood were flushed. Vessels were infected with *K. pneumoniae* intraluminally for 1 hour prior to fixation in 4% paraformaldehyde solution (PFA) for 1 hour. Fixed tissue was then washed thoroughly with PBS before being placed in 10% sucrose solution in PBS for 24 hours and a further 24 hours in 30% sucrose solution.

For cryosectioning, samples were embedded in OCT matrix and cut to produce 5 µm sections. Standard protocols were used for haematoxylin and eosin (H&E) staining of the sections, which were then imaged on an Eclipse 80i microscope with a 40x objective, using the NIS-Elements acquisition programme by Nikon. Scoring was carried out in a blinded manner by one researcher from the laboratory who was not involved in this research. The scoring criteria are detailed in Table S3.

Six vessels from five different rats were analyzed.

### RNA isolation and RT-qPCR

RNA was isolated and extracted by lysing cells in TRIzol reagent using the manufacturer provided guidance. RNA concentration and purity was determined using a Nanodrop spectrophotometer by assessing the A260/A280 ratio. Duplicate cDNA preparations from each sample were generated from 1 μg of RNA using Moloney murine leukaemia virus (M-MLV) reverse transcriptase (Sigma-Aldrich) according to the manufacturer’s instructions. Quantitative real-time PCR analysis of gene expression was undertaken using the KAPA SYBR FAST qPCR Kit and a Rotor-Gene Q real-time PCR cycler System (Qiagen). Thermal cycling conditions were as follows: 95 °C for 3 min for enzyme activation, 40 cycles of denaturation at 95 °C for 10 s and annealing at 60 °C for 20 s. Primers used in qPCR reactions are listed in Table S2. cDNA samples were tested in duplicate and relative mRNA quantity was determined by the comparative threshold cycle (ΔΔCT) method, using glyceraldehyde 3-phosphate dehydrogenase (hGAPDH) gene normalisation.

### Statistical analysis

Data are represented as mean with SD. Two groups of data were compared using an unpaired, two-tailed student’s t-test. Multiple groups were compared using a one-way or two-way ANOVA followed by Tukey’s or Dunnet’s multiple group comparison correction, respectively. For concentration response curves, curves were fit using log (agonist) vs response (3 parameters) approach and statistical significance determined using two-way ANOVA, including Dunnet’s multiple group comparison correction. Analyses were performed using Prism software (GraphPad, version 9.02).

## ACKNOWLEDGEMENTS

We thank the members of the J.A.B. and T.M.C. laboratories for their thoughtful discussions and support with this project. S.R. and C. R. are the recipients of PhD fellowships funded by the Department for Employment and Learning (Northern Ireland, UK). This work was supported by Biotechnology and Biological Sciences Research Council (BBSRC) (BB/V007939/1) and Medical Research Council (MRC) (MR/V032496/1) funds to J.A.B; Health & Social Care R&D Division, Northern Ireland (STL/4748/13), Medical Research Council (MC_PC_15026), and Queen’s University Belfast Agility Fund funds to T.M.C; and British Heart Foundation programme grant (RG/F/20/110007) to J. McC.

## DECLARATION OF INTERESTS

J.A.B. declares consultancy fees from VaxDyn and GSK. The other authors have declared that no conflict of interest exists.

## SUPPLEMENTARY FIGURE LEGENDS

Figure S1. *K. pneumoniae* inhibits ACh-induced vasodilation.

(A). Representative trace of the effect of NO pathway inhibitor L-NAME on ACh-induced vasodilation in non-infected PE-constricted vessels.

(B). Representative trace of the effect of EDH pathway inhibitors TRAM-34 and apamin on ACh-induced vasodilation in non-infected PE-constricted vessels.

(C). Representative trace of the effect of NO and EDH pathway inhibitors L-NAME, TRAM-34 and apamin on ACh-induced vasodilation in non-infected PE-constricted vessels.

(D). Outer diameter of non-infected and infected vessels for 1 h with Kp43816.

(E). Outer diameter of the non-infected and infected vessels for 1 h with Kp43816 after PE-induced vasoconstriction.

(F). Concentration response of ACh-induced peak dilation in non-infected and infected vessels with 10^4^ and 10^3^ CFUs of strain Kp43816.

(G). Representative trace of Kp43816 (10^3^ CFUs)-triggered inhibition of ACh-induced vasodilation in PE-constricted vessels.

(H). Peak dilation of non-infected and infected vessels treated with the NO donor SNP in PE-constricted vessels.

(I). Dose response of ACh-induced peak dilation in non-infected and infected vessels with UV-killed Kp43186 and 10^4^ latex beads.

In A, B, C, and G the traces are representative of six vessels from three rats. In F, and I peak dilation is shown as mean ± SD from six vessels from three rats per group. In D and H, dilation is shown as mean ± SD from six and four vessels, respectively, from three rats per group. In E, constriction is shown as mean ± SD from six vessels from three rats per group. In F and I, data were analysed with two-way ANOVA and Dunnet’s multiple group comparison correction. The asterisks indicate p <0.0001 (****). In D, E and H, data was analysed using an unpaired, two-tailed student’s t-test; ns, not significant.

Figure S2. T6ss effector VgrG4 inhibits ACh-induced vasodilation.

(A). Representative trace of the lack of effect of infection with the *cps* mutant (Δ*manC*; 43816-Δ*manC*) on ACh-induced vasodilation in PE-constricted vessels.

(B). Representative trace of the lack of effect of infection with the *tssB* mutant (Δ*tssB*; 43816-Δ*tssB*) on ACh-induced vasodilation in PE-constricted vessels.

(C). Concentration response of ACh-induced peak dilation in non-infected and infected vessels with strains NTUH-K2044 and the T6SS mutant NTUH-Δ*clpV*.

(D). Representative trace of the lack of effect of infection with the *vgrG4* mutant (Δ*vgrG4*; 43816-Δ*vgrG4*) on ACh-induced vasodilation in PE-constricted vessels.

(E). Representative trace of the lack of effect of infection with Ye on ACh-induced vasodilation in PE-constricted vessels.

(F). Representative trace of the effect of infection with YeVgrG4 on ACh-induced vasodilation in PE-constricted vessels.

In A, B, D, E and F the traces are representative of six vessels from three rats. In F, peak dilation is shown as mean ± SD from six vessels from three rats per group and data were analysed with two-way ANOVA and Dunnet’s multiple group comparison correction. The asterisks indicate p <0.001 (***).

Figure S3. *K. pneumoniae* infection and Ca2+ signalling in endothelium.

(A). Concentration response of ACh-induced intracellular Ca^2+^ in non-infected HULEC cells.

(B). 4-DAMP, M3 muscarinic receptor antagonist, blunts average ACh-induced Ca^2+^ signalling in non-infected and Kp43816-infected vessels.

In A, data is shown as mean ± SD from three independent experiments in sextuplicate. In B, each dot represents the data from one vessel before and after infection.

Figure S4. *K. pneumoniae* targets the phosphorylation of eNOS Ser^1177^ and Thr^495^.

(A). Immunoblot analysis of eNOS and tubulin levels in infected HPMEC cells for the indicated time points. Densitometry analysis of blots (n = 3) representing the ratio of eNOS versus tubulin.

(B). Immunoblot analysis of phosphorylated eNOS Ser^1177^, eNOS and tubulin levels in HULEC cells treated with condition media from control medium containing 1 μM bradykinin (Bk), histamine (His), and ionomycin (Iono), and media in which the agonists were incubated with Kp43816 for 5 min, and then bacteria were spun down. Densitometry analysis of blots (n = 3) representing the ratio of phosphorylated eNOS Ser^1177^ versus eNOS.

(C). Immunoblot analysis of phosphorylated eNOS Thr^495^ and tubulin levels in infected HPMEC cells for the indicated time points. Densitometry analysis of blots (n = 3) representing the ratio of phosphorylated eNOS Thr^495^ versus eNOS.

(D). Immunoblot analysis of phosphorylated eNOS Thr^495^, eNOS and tubulin levels in non-infected and infected HULEC cells treated for 2 min with 1 μM bradykinin (Bk), histamine (His), and ionomycin (Iono). Densitometry analysis of blots (n = 3) representing the ratio of phosphorylated eNOS Thr^495^ versus eNOS.

(E). Immunoblot analysis of phosphorylated eNOS Thr^495^, eNOS and tubulin levels in non-infected and infected HULEC cells with strains Kp43816 and the the T6SS *tssB* (Δ*tssB*; 43816-Δ*tssB*) for the indicated time points. Densitometry analysis of blots (n = 3) representing the ratio of phosphorylated eNOS Thr^495^ versus eNOS.

Blots are representative of three independent experiments. In all panels, data were compared against the non-infected control using one-way ANOVA and Tukey’s multiple group comparison correction. The asterisks indicate p < 0.05 (*), <0.01 (**). <0.001 (***), or <0.0001 (****). ns, not significant.

Figure S5. *K. pneumoniae* induces PP2Ac phosphatase to target eNOS Ser^1177^.

(A). Immunoblot analysis of PP2Ac and tubulin levels in infected HPMEC cells for the indicated time points. Densitometry analysis of blots (n = 3) representing the ratio of PP2Ac versus tubulin.

(B). Immunoblot analysis of PP2AC and tubulin levels in cells transfected with All Stars siRNA control, or PP2Ac siRNA. Densitometry analysis of blots (n = 3) representing the percentage of PP2Ac reduction in transfected cells with All Stars siRNA control, normalized to 100%, or PP2Ac siRNA.

(C). Immunoblot analysis of phosphorylated EGFR and tubulin levels in HULEC cells transfected with All Stars siRNA control, or *tlr4* siRNA. And infected with Kp43816 for the indicated time points. Densitometry analysis of blots (n = 3) representing the ratio of PP2Ac versus tubulin.

(D). Immunoblot analysis of TLR4 and tubulin Densitometry analysis of blots (n = 3) representing the percentage of TLR4 reduction in transfected cells with All Stars siRNA control, normalized to 100%,

(E). Immunoblot analysis of phosphorylated AKT, PAK4, ERK and GSK3β, and tubulin levels in HUELC cells infected with Kp43816 for the indicated time points. Densitometry analysis of blots (n = 3) representing the ratio of phosphorylated protein versus tubulin.

Blots are representative of three independent experiments. In all panels, data were compared against the non-infected control using one-way ANOVA and Tukey’s multiple group comparison correction. The asterisks indicate p < 0.05 (*), <0.01 (**). <0.001 (***) or. ns, not significant.

Figure S6. *K. pneumoniae* activates PKCβ kinase to phosphorylate eNOS Thr^495^.

(A). Immunoblot analysis of phosphorylated PKC and tubulin levels in infected HPMEC cells for the indicated time points. Densitometry analysis of blots (n = 3) representing the ratio of phosphorylated PKC versus tubulin.

(B). Immunoblot analysis of phosphorylated eNOS Thr^495^, eNOS and tubulin levels in infected HULEC cells treated with vehicle solution DMSO and the PKC inhibitor RO 31-8220 for the indicated time points. Densitometry analysis of blots (n = 3) representing the ratio of phosphorylated eNOS Thr^495^ versus eNOS.

(C). Immunoblot analysis of phosphorylated eNOS Thr^495^, eNOS and tubulin levels in HULEC cells transfected with All Stars siRNA control, PKCα or PKCε siRNA infected with Kp43816 for the indicated time points. Densitometry analysis of blots (n = 3) representing the ratio of phosphorylated eNOS Thr^495^ versus eNOS.

(D). Efficiency of transfection of siRNA into HULEC cells. Efficiency of transfection presented as percent knockdown after transfection. mRNA levels of the indicated PKC transcripts were accessed 48 h post transfection as fold change against control non silencing agents (AS [AllStars control, non silencing siRNA] after gene normalization. Values are presented as the means ± SD from three independent experiments measured in duplicates.

(E). Immunoblot analysis of phosphorylated eNOS Thr^495^, eNOS and tubulin levels in infected HULEC cells and treated with vehicle solution DMSO and the PKCβ inhibitor LY333-531 for the indicated time points. Densitometry analysis of blots (n = 3) representing the ratio of phosphorylated eNOS Thr^495^ versus eNOS.

(F) Efficiency of transfection presented as percent knockdown after transfection. mRNA levels of the NLRX1 transcripts were accessed 48 h post transfection as fold change against control non silencing agents (AS [AllStars control, non silencing siRNA] after gene normalization. Values are presented as the means ± SD from three independent experiments measured in duplicates.

Figure S7. Treatment with MitoTEMPO and PKCβ inhibitor restores ACh-induced vasodilation in *K. pneumoniae*-infected vessels.

(A). Representative trace of the lack of effect of MitoTEMPo on ACh-induced vasodilation in non-infected PE-constricted vessels.

(B). Representative trace of the effect of MitoTEMPO on ACh-induced vasodilation in Kp43816-infected PE-constricted vessels.

(C). Representative trace of the effect of MitoTEMPO and L-NAME on ACh-induced vasodilation in Kp43816-infected PE-constricted vessels.

(D). Representative trace of the effect of MitoTEMPO, TRAM-34 and apamin on ACh-induced vasodilation in Kp43816-infected PE-constricted vessels.

(E). Representative trace of the lack of effect of the PKCβ inhibitor LY333-531 on ACh-induced vasodilation in non-infected PE-constricted vessels.

(F). Representative trace of the effect of LY333-531 on ACh-induced vasodilation in Kp43816-infected PE-constricted vessels.

(G). Representative trace of the effect of LY333-531 and L-NAME on ACh-induced vasodilation in Kp43816-infected PE-constricted vessels.

(H). Representative trace of the effect of LY333-531 on ACh-induced vasodilation in PE-constricted vessels infected with the *cps* mutant (Δ*manC*; 43816-Δ*manC*).

(I). Representative trace of the effect of LY333-531 and L-NAME on ACh-induced vasodilation in PE-constricted vessels infected with the *cps* mutant (Δ*manC*; 43816-Δ*manC*).

(J). Representative trace of the effect of LY333-531, TRAM-34 and apamin on ACh-induced vasodilation in PE-constricted vessels infected with the *cps* mutant (Δ*manC*; 43816-Δ*manC*).

(K). Representative trace of the effect of LY333-531, L-NAME, TRAM-34 and apamin on ACh-induced vasodilation in PE-constricted vessels infected with the *cps* mutant (Δ*manC*; 43816-Δ*manC*).

The traces are representative of six vessels from three rats.

Figure S8. *K. pneumoniae* disrupts vasodilation by targeting eNOS post translational modifications.

Working model of the strategies deployed by *Klebsiella* to abrogate ACh-induced vasodilation by affecting the NO and EDH pathways controlling vasodilation in the vessels probed in this work. *Klebsiella* targets eNOS post translational modifications to blunt the NO pathway. *Klebsiella* activates a TLR4-EGFR-dependent pathway to increase the phosphatase PP2Ac to abrogate the phosphorylation of the eNOS Ser^1177^ active site whereas the T6SS effector VgrG4 via the innate receptor NLRX1 induces mtROS which licences the PKCβ kinase to phosphorylate eNOS inhibitory site Thr^495^. VgrG4-induced mtROS attenuates the EDH pathway.

Figure S9. Effect of inhibitors on *K. pneumoniae* growth.

(A). Growth of Kp43186 in LB in the presence of vehicle solution DMSO, and the inhibitors Ro 31-8220 and LY333-531.

(B). Growth of Kp43186 in M9 minimal medium in the presence of vehicle solution DMSO, and the inhibitors Ro 31-8220 and LY333-531.

In A and B, results are shown as mean ± SD from four independent experiments.

