## Supplementary figures and images for "*Klebsiella pneumoniae* disrupts vasodilation by targeting eNOS post translational modifications via the type VI secretion system and the capsule polysaccharide"

### Supplementary figures

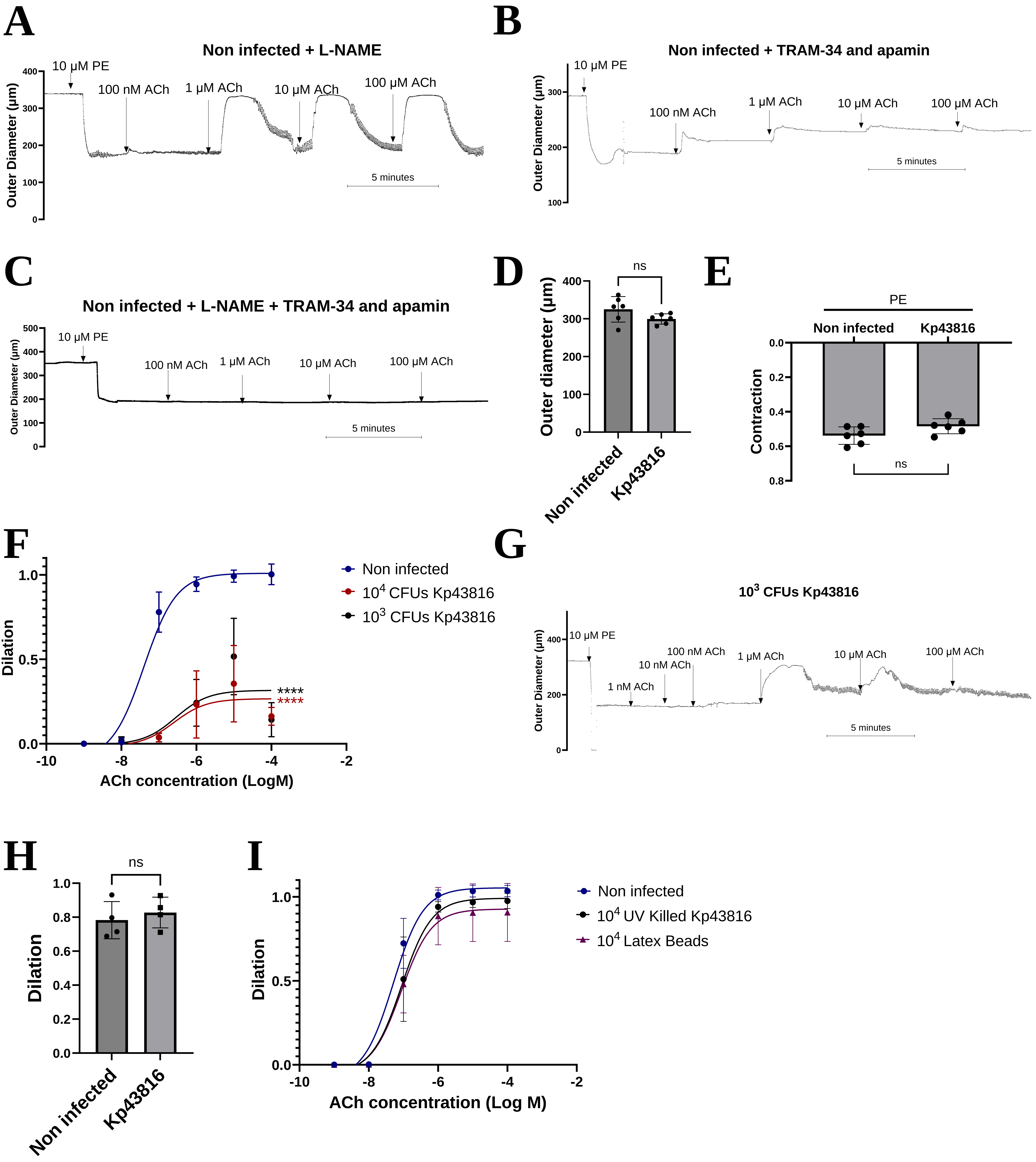

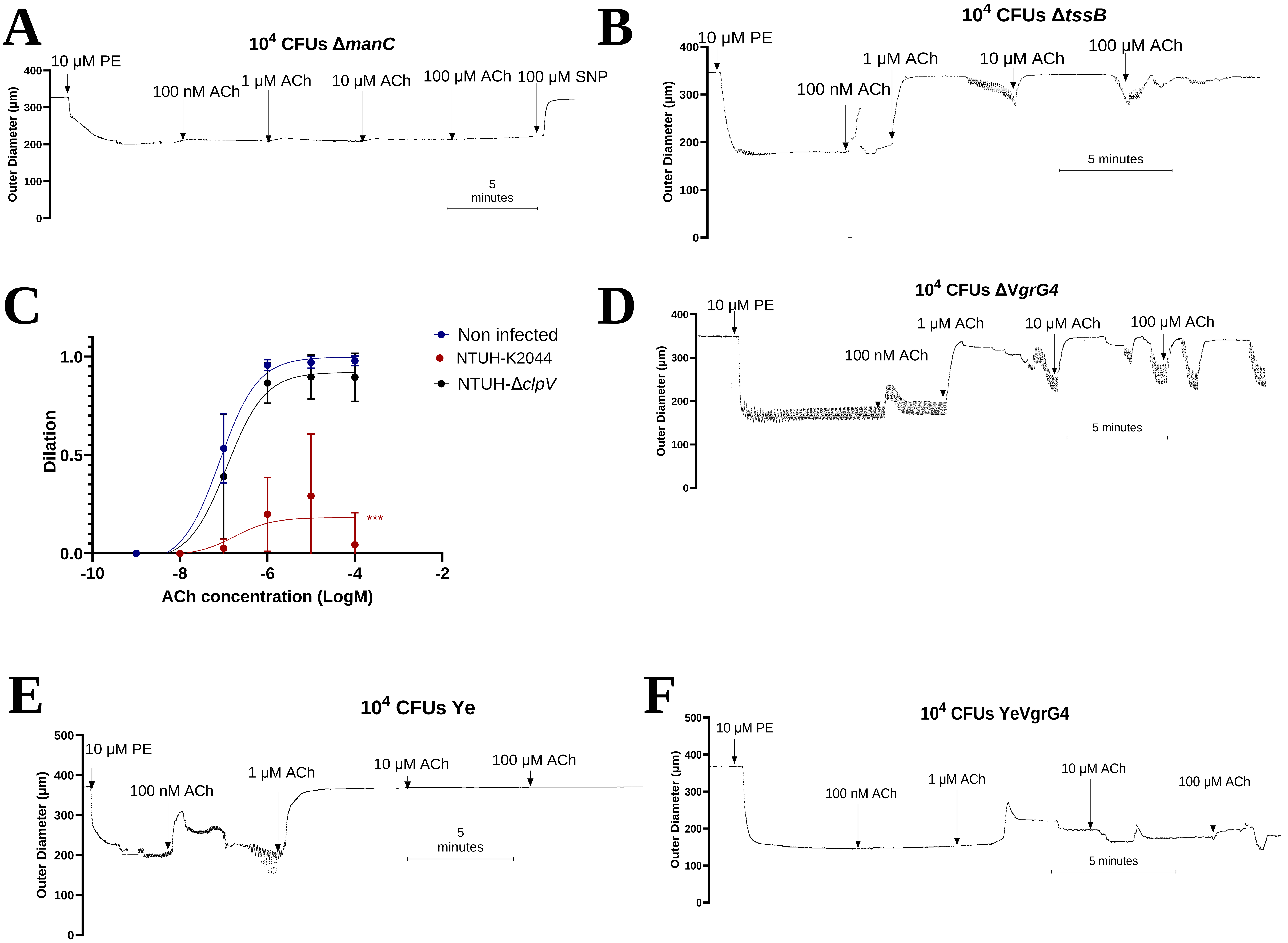

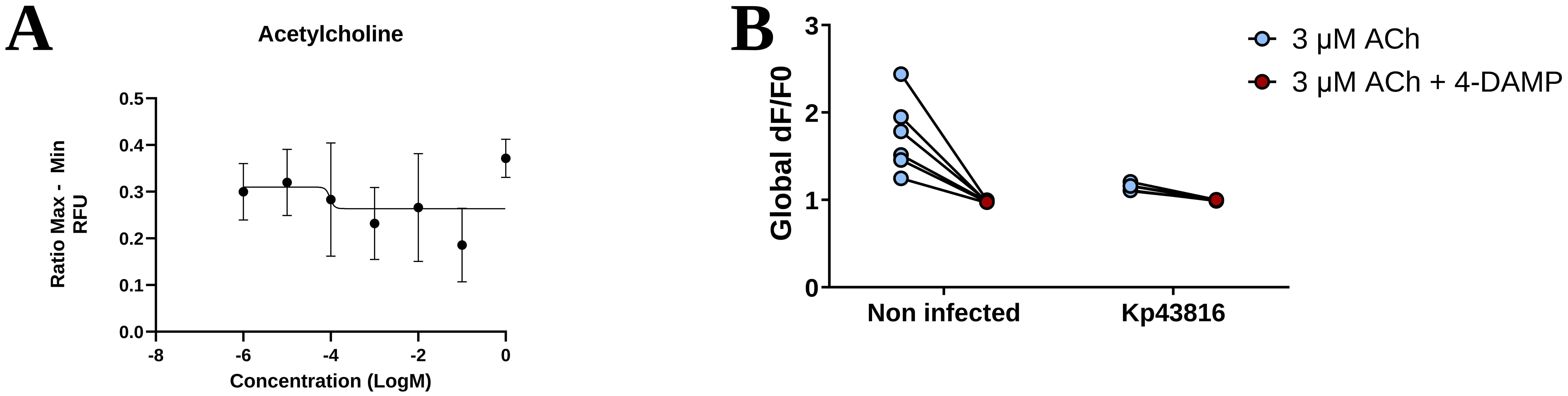

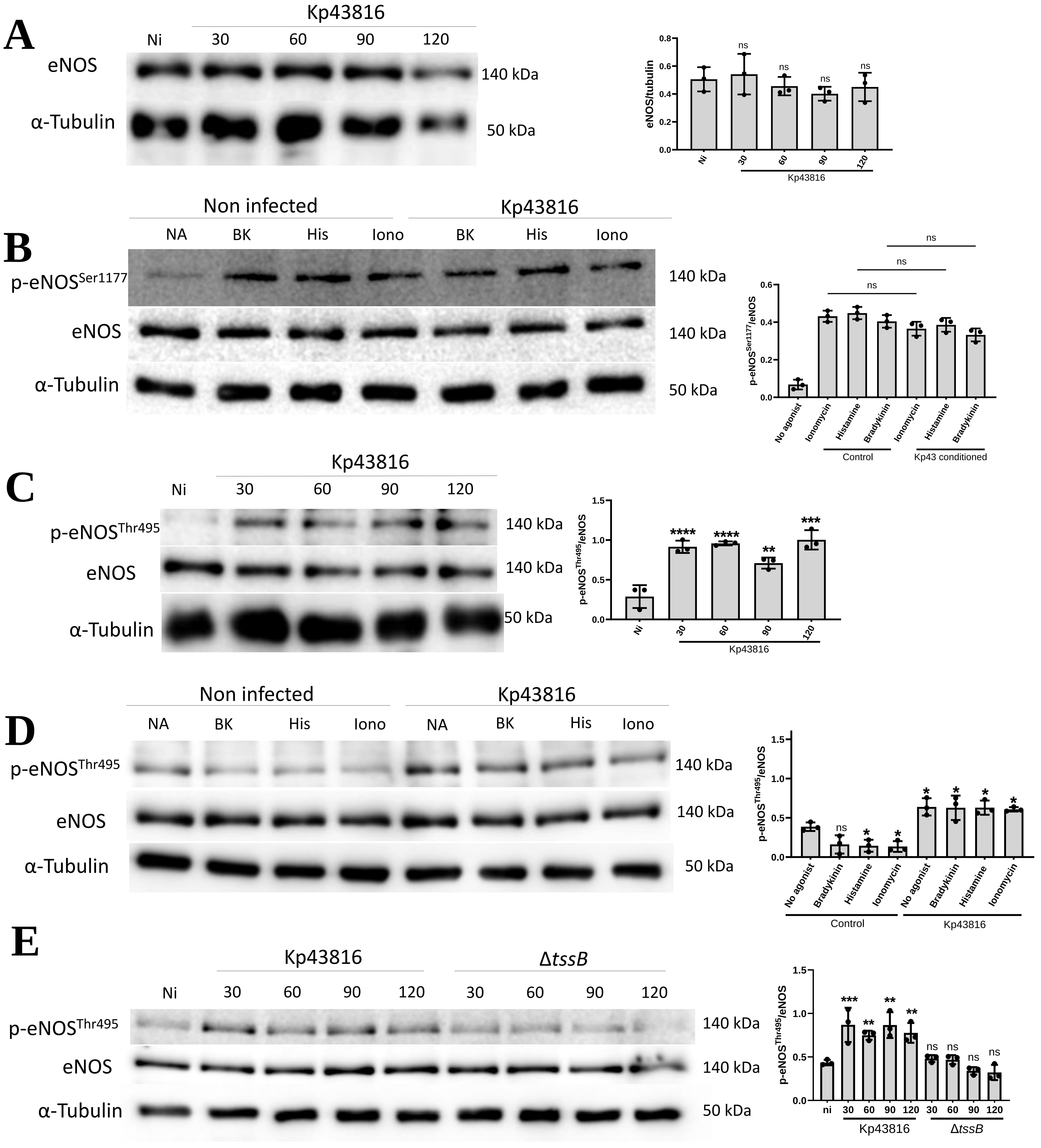

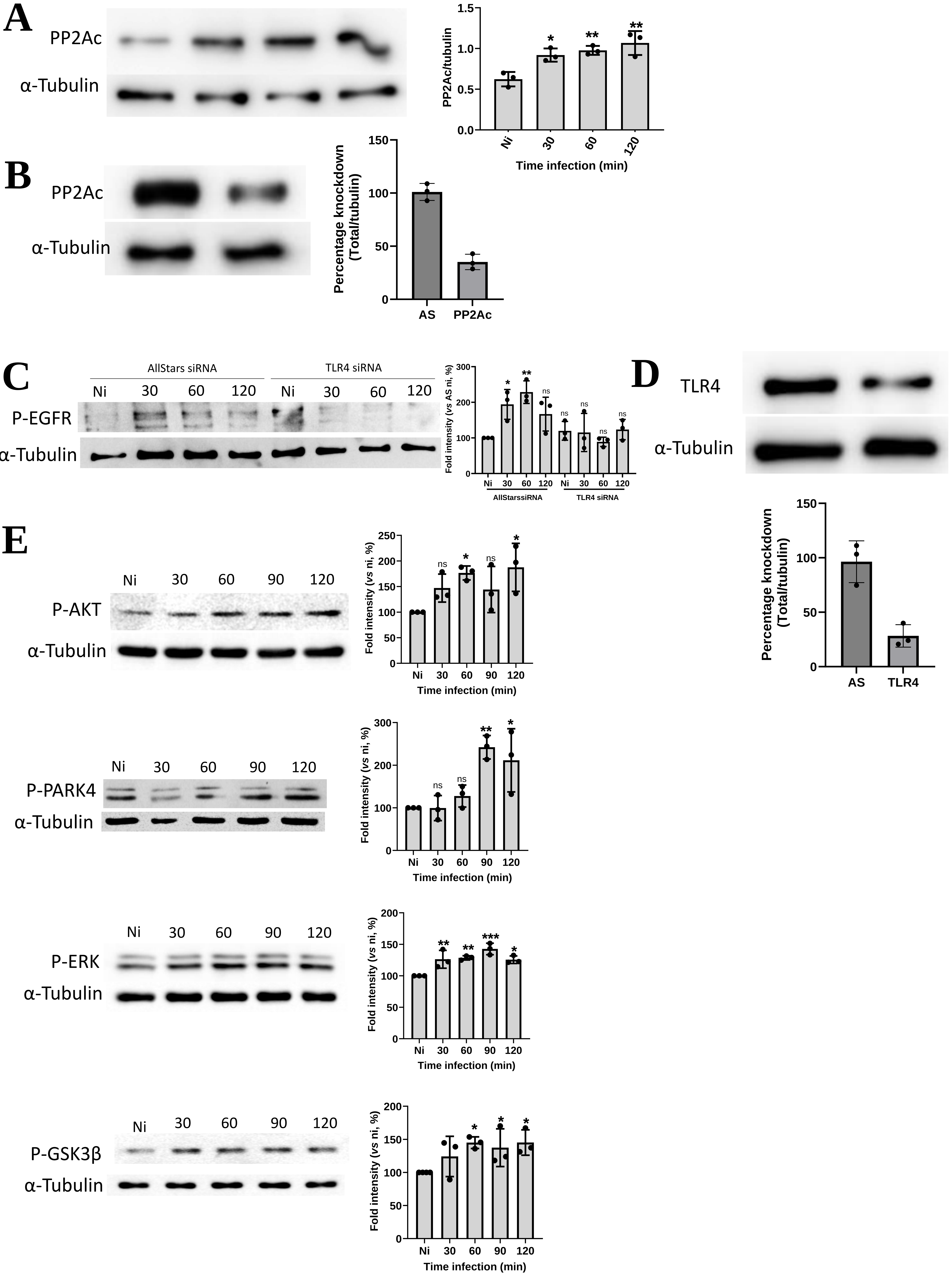

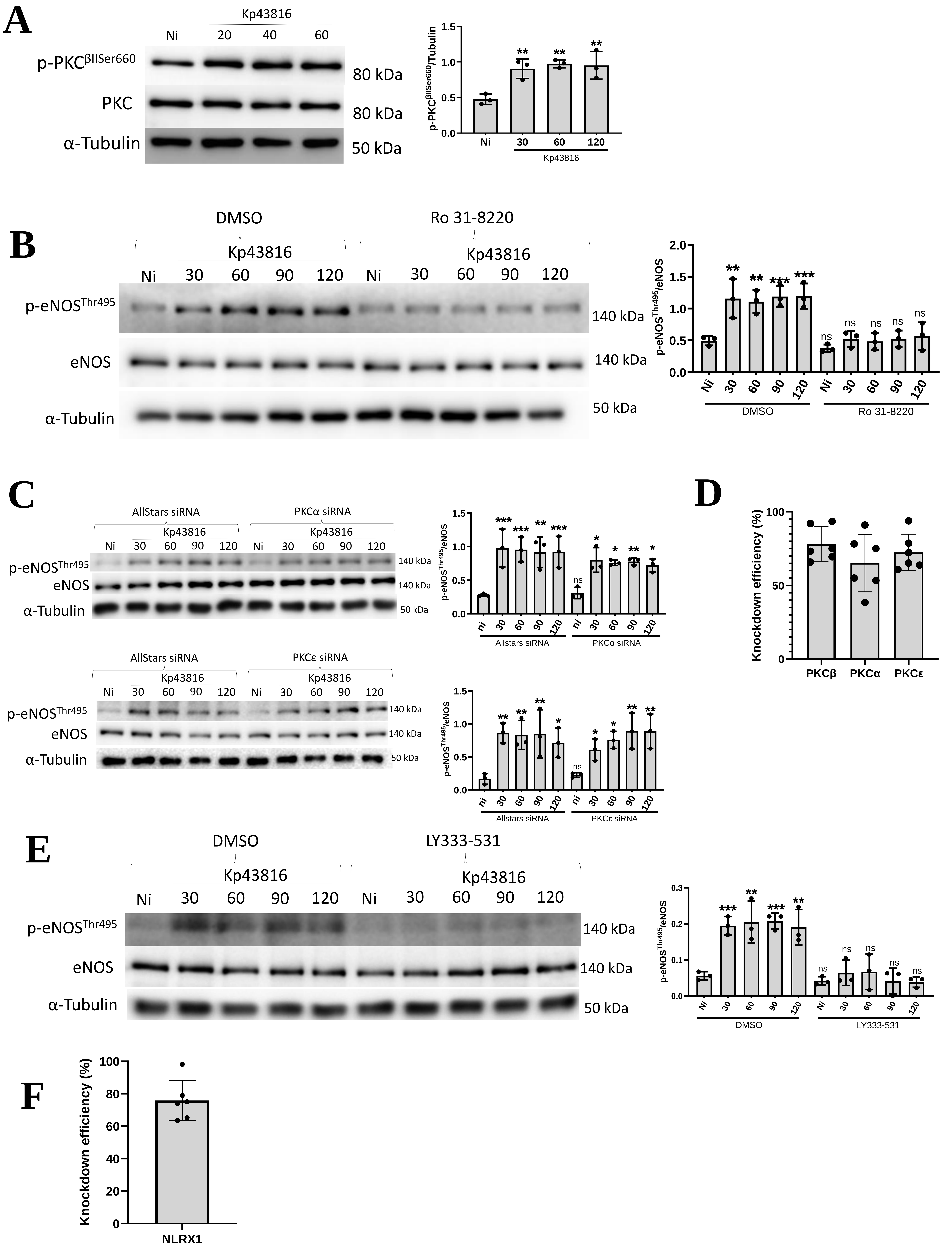

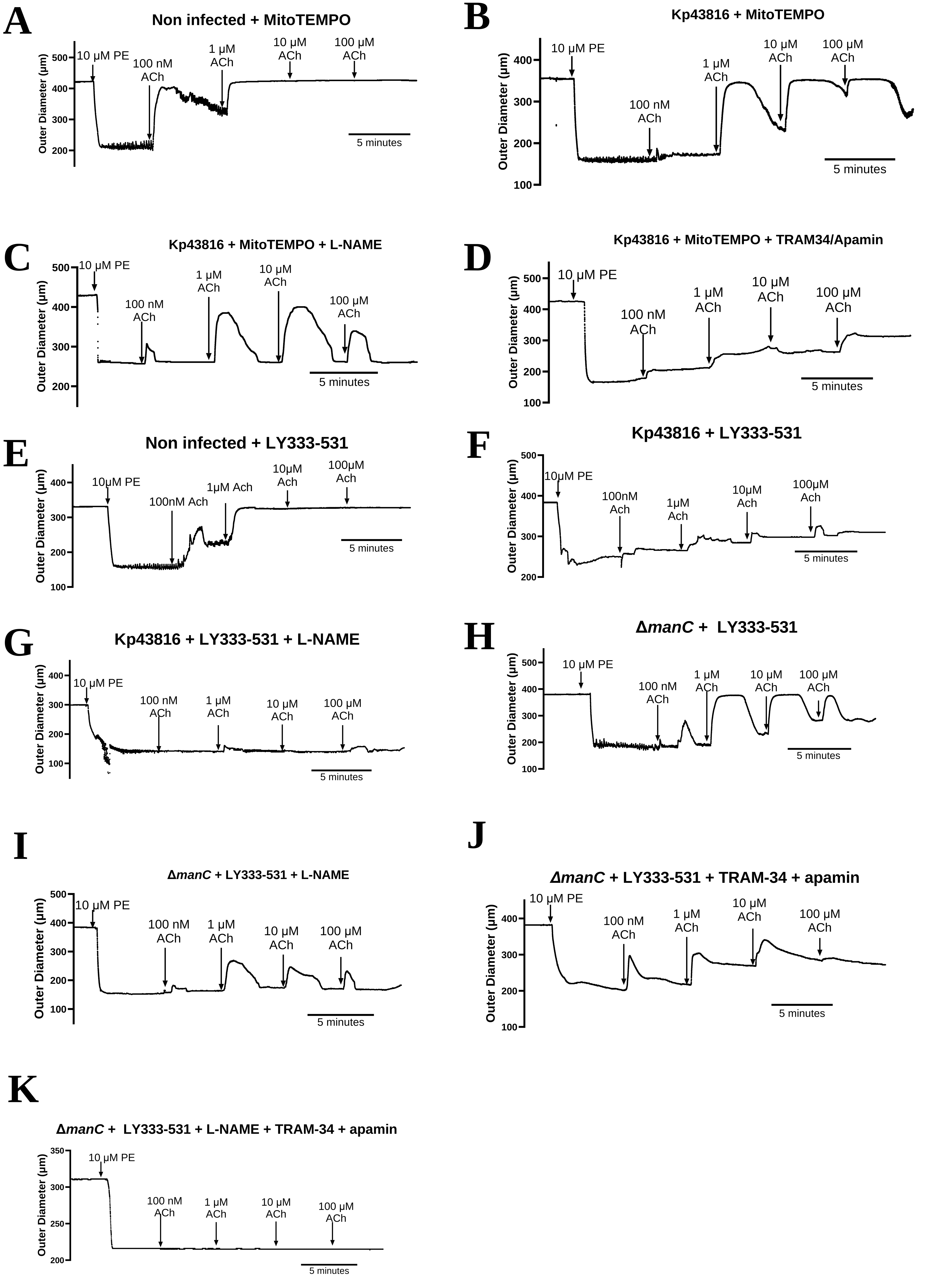

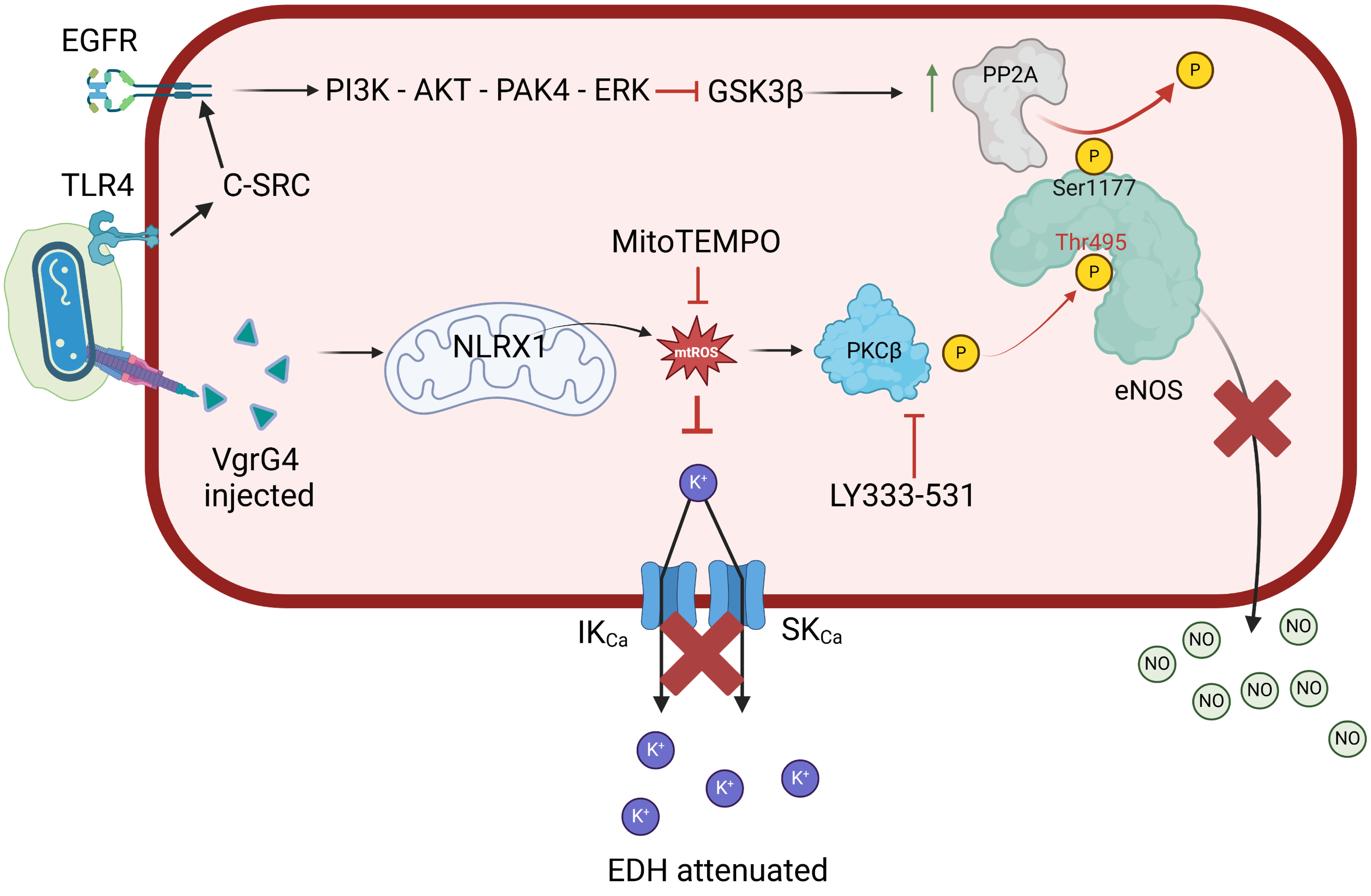

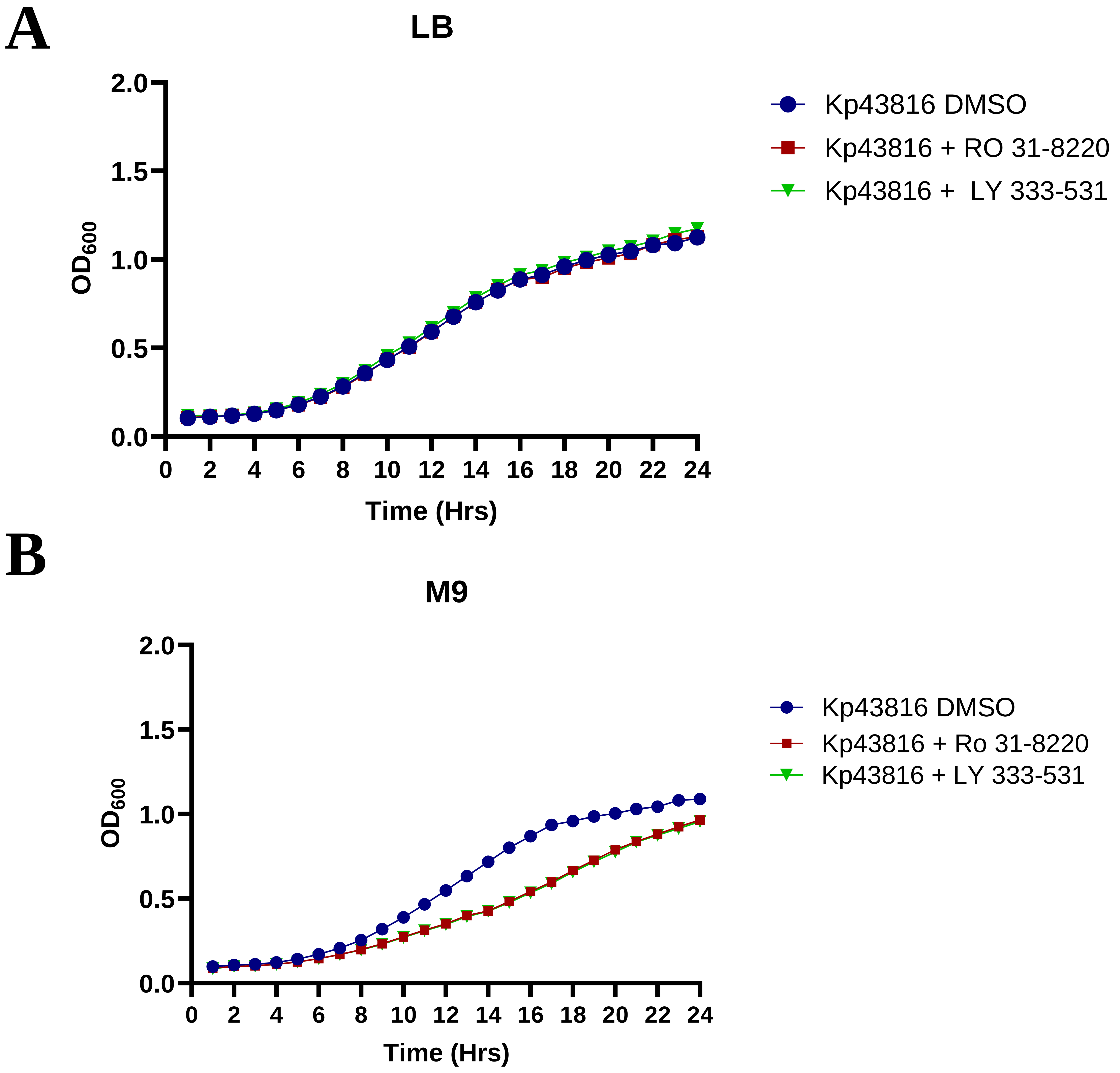
