## Supplementary material for "*Klebsiella pneumoniae* disrupts vasodilation by targeting eNOS post translational modifications via the type VI secretion system and the capsule polysaccharide": Table S1

**Table S1. Bacterial strains used in this work.**

| **Strain** | **Genotype or comments.** | **Reference** |
| --- | --- | --- |
| *Klebsiella pneumoniae* |  |  |
| NTUH-K2044 | Clinical isolate; serotype O1:K2; sequence type ST23 | 1 |
| ATCC43816 | Clinical isolate; serotype O1:K2; sequence type ST493 | ATCC |
| MRSN-14444 | Clinical isolate; MRSN Diversity Panel; sequence type 4270 | 2 |
| AKP5 | Clinical isolate; carbapenem resistant | Karolinska Institute collection |
| 43Δ*manCKm* | ATCC43816, Δ*manCKm*, *manC* gene inactivated, Km^R^ | 3 |
| 43816-Δ*tssB* | ATCC43816, Δ*tssB, tssB* gene inactivated | 4 |
| 43816-Δ*vgrG4* | ATCC43816, Δ*vgrG4, vgrG4* gene inactivated | This work |
| NTUH-Δ*clpV* | NTUH-K2044, ΔclpV*, clpV* gene inactivated | 4 |
| *Yersinia enterocolitica* |  |  |
| Ye | pYV negative derivative of strain WA-314 serotype O:8, harbouring the pT3SS plasmids; Spec^R^ | 5 |
| YeVgrG4 | Ye strain harbouring the plasmids pT3SS and pE_53_-*vgrG4* with a VSVG tagged *vgrG4*; Spec^R^ Cm^R^ | 6 |
| *Escherichia coli* |  |  |
| JKE201 | MFDpirΔ*mcrA*Δ(*mrr*-*hsd*RMS-*mcr*BC) *aac(3)3IV::lacI^q^* | 7 |

6.	Sá-Pessoa, J., López-Montesino, S., Przybyszewska, K., Rodríguez-Escudero, I., Marshall, H., Ova, A., Schroeder, G.N., Barabas, P., Molina, M., and Curtis, T. (2023). A trans-kingdom T6SS effector induces the fragmentation of the mitochondrial network and activates innate immune receptor NLRX1 to promote infection. Nature Communications *14*, 871.

7.	Harms, A., Liesch, M., Körner, J., Québatte, M., Engel, P., and Dehio, C. (2017). A bacterial toxin-antitoxin module is the origin of inter-bacterial and inter-kingdom effectors of Bartonella. PLoS genetics *13*, e1007077.
