## Supplementary material for "*Klebsiella pneumoniae* disrupts vasodilation by targeting eNOS post translational modifications via the type VI secretion system and the capsule polysaccharide": Table S2

**Table S2. Primers used in this study.**

| **Name** | **Forward (5’-3’)** | **Reverse (5’-3’)** | **Purpose** |
| --- | --- | --- | --- |
| h_PRKCA | AGAACGTGCACGAGGTGAA | ACCCACAGTGATCGCAGAAG | siRNA knockdown efficiency |
| h_PRKCB | GGCAGA AGA ACGTGCATGAG | CAA AACGTGGGGCTGGAG TA | siRNA knockdown efficiency |
| h_PRKCE | ACAAAATCACCAACAGCGGC | GTTGAACTCATCCAGGCCCA | siRNA knockdown efficiency |
| h_NLRX1 | GTGCCCGGAAGCTGGGCTTG | CCGGGCACCACCTTCAGCAG | siRNA knockdown efficiency |
| h_GAPDH | GAGAAGGCTGGGGCTCATTT | AGTGATGGCATGGACTGTGG | siRNA knockdown efficiency |
| VgrG4UP | GAATTCGCAACGATAAATCCAATCAA | GGATCCTATATTCCTGTTTTACTCCG | VgrG4 mutagenesis |
| VgrG4DOWN | GGATCCACAATGATA TTGTTACCCCG | GAATTCCGTTTATTC TCCTGCGGC | VgrG4 mutagenesis |
| VgrG4SCREEN | AACTTCCTCCGCTGAACCTG | TGCTTTGGCAAAATTTTCGTGC | VgrG4 mutagenesis |
