## Supplementary material for "*Klebsiella pneumoniae* disrupts vasodilation by targeting eNOS post translational modifications via the type VI secretion system and the capsule polysaccharide": Table S3

**Table S3. Histology scoring criteria.**

| **Damage** | **Score** |
| --- | --- |
| Major disruption in endothelium | 5 |
| Major disruption in vascular smooth muscle layer | 5 |
| Minor disruption in endothelium | 2 |
| Minor disruption in vascular smooth muscle layer | 2 |
| Small breaks in endothelium | 1 |
| Small breaks in vascular smooth muscle layer | 1 |
